# Comprehensive annotation of 3′UTRs from primary cells and their quantification from scRNA-seq data

**DOI:** 10.1101/2021.11.22.469635

**Authors:** Mervin M. Fansler, Sibylle Mitschka, Christine Mayr

**Affiliations:** Tri-Institutional Training Program in Computational Biology and Medicine, Weill-Cornell Graduate College, New York, NY 10021, USA; Cancer Biology and Genetics Program, Memorial Sloan Kettering Cancer Center, New York, NY 10065, USA

## Abstract

Approximately half of human genes generate mRNA isoforms that differ in their 3′UTRs while encoding the same protein. 3′UTR and mRNA length is determined by 3′ end cleavage sites (CS). Here, we mapped and categorized mRNA 3′ end CS in more than 200 primary human and mouse cell types, resulting in a 40% increase of CS annotations relative to the GENCODE database. We incorporated these annotations into a novel computational pipeline, called scUTRquant, for rapid, precise, and accurate quantification of gene and 3′UTR isoform expression from single-cell RNA sequencing (scRNA-seq) data. When applying scUTRquant to data from 474 cell types and 2,134 perturbations, we discovered extensive 3′UTR length changes across cell types that are as widespread and dynamically regulated as gene expression changes. Our data indicate that mRNA abundance and mRNA length are two independent axes of gene regulation that together determine the amount and spatial organization of protein synthesis.

## Introduction

In mRNAs, the 3′ untranslated region (3′UTR) is located between the stop codon of the coding sequence and the poly(A) tail. Pre-mRNA cleavage and polyadenylation (CPA) is initiated upon recognition of the polyadenylation signal (PAS) by the CPA machinery and determines mRNA and 3′UTR length^1^. Approximately half of human genes use alternative cleavage and polyadenylation (APA) to generate mRNA isoforms that encode the same protein but that differ in their 3′UTRs^2^. In addition, ∼25% of genes use intronic polyadenylation (IPA) signals to generate mRNA isoforms with alternative last exons, thus producing different protein isoforms^3-7^. APA is tightly controlled during development and dysregulated in disease^8,9^. Alterations in 3′UTR length affect the presence of binding sites for microRNAs and RNA-binding proteins and can regulate mRNA stability and translation^10,11^. More recently, 3′UTRs have emerged as important regulators of subcytoplasmic location of translation and mRNA-dependent co-translational protein complex assembly^12-21^, reviewed in^1,22^.

When APA was initially discovered to be a widespread phenomenon, it was thought to be a new mode of gene expression regulation, where a switch in 3′UTR isoform usage changes overall gene expression^10,11^. However, several labs that had performed transcriptome-wide APA analyses found that fewer than 20% of 3′UTR changes regulate mRNA or protein abundance.

This suggested that mRNA abundance and 3′UTR length may be independent gene outputs^2,23-27^. These analyses were performed on a limited set of cell types, and it remains unclear whether the independence of gene and 3′UTR expression is a general phenomenon.

Whereas gene expression analysis has been performed for nearly 30 years^28,29^, differential 3′UTR analysis is a more recent development that initially required the use of custom 3′ end sequencing protocols^2,3,24^, which limited its widespread use. Now, 3′-tag-based single-cell RNA sequencing (scRNA-seq) protocols, such as 10x Genomics, can be used to quantify differential 3′UTRs^30-34^. However, only few reads from 10x Genomics data cross mRNA 3′ end cleavage sites (CS), which makes CS identification difficult. Therefore, current APA pipelines either use *de novo* peak calling without CS identification^30,31^ or they assign reads to the closest known CS, obtained from CS databases^32,34-36^. When using *de novo* peak calling methods, variations in sequencing depth and batch effects usually limit comparability across studies. In contrast, the reliance on reference CS yields more consistent results, but the results strongly depend on the quality of the reference annotation. As existing 3′ end CS annotations, such as PolyA_DB V3.2 and PolyASite 2.0, were predominantly generated from cell line data, CS found only in rare cell types may be missing^35,36^.

To provide a fast and reliable workflow for simultaneous gene expression and 3′UTR quantification from scRNA-seq data, we developed a quantification pipeline, called scUTRquant and a statistical testing package, called scUTRboot. To quantify 3′UTRs with precise 3′ ends, we first generated a comprehensive CS annotation based on Microwell-seq (MWS) data of 206 human and mouse cell types^37,38^. Our new MWS CS annotation was generated from 7 billion CS-spanning reads and expands GENCODE CS annotations by 40%. To demonstrate potential applications of our new analysis workflow, we applied it to 474 cell types and 2,134 perturbations and observed that changes in gene expression and in 3′UTR length occurred in different groups of genes, indicating that they largely represent independent regulatory events. Our large-scale analysis revealed that only about half of gene regulatory events cause changes in mRNA abundance greater than 1.5 fold, whereas the other half affects mRNA length, which may impact the spatial control of protein synthesis.

## Results

### Mapping and characterization of mRNA 3′ end CS in 206 primary cell types

Reads obtained from 10x Genomics rarely contain untemplated adenosines, but approximately 35% of reads obtained from MWS span mRNA 3′ end CS (Fig. 1a)^37,38^. MWS was previously applied to 104 and 102 primary cell types from mouse and human, and we were able to use 7 billion CS-spanning reads to map mRNA 3′ ends at single-nucleotide resolution (Fig. 1b). Briefly, we removed poly(A) tails and mapped the remaining portions of the reads to the genome, followed by filtering of low abundance reads and peaks derived from priming at genomic adenosine stretches. The resulting mRNA 3′ end CS were intersected with GENCODE annotations and classified into three groups, (i) common CS (previously annotated in GENCODE and observed in MWS data), (ii) MWS-only CS (not previously annotated), and (iii) GENCODE-only CS (annotated in GENCODE but not detected in MWS data). We include GENCODE annotations in our new MWS CS annotation, which expands the number of CS present in GENCODE annotations by 40% (Fig. 1b). Two thirds of the novel sites were detected in fewer than 10 cell types, suggesting that their absence in current annotations may indeed be caused by their cell type-specific expression patterns (Fig. S1a). Novel CS were derived from tissues that span all developmental stages, including embryo, fetal, neonatal, and adult tissues. The largest fractions of novel CS were obtained from adult tissues, such as omentum, pleura, and testis (Fig. S1b, S1c).

**Figure 1.**
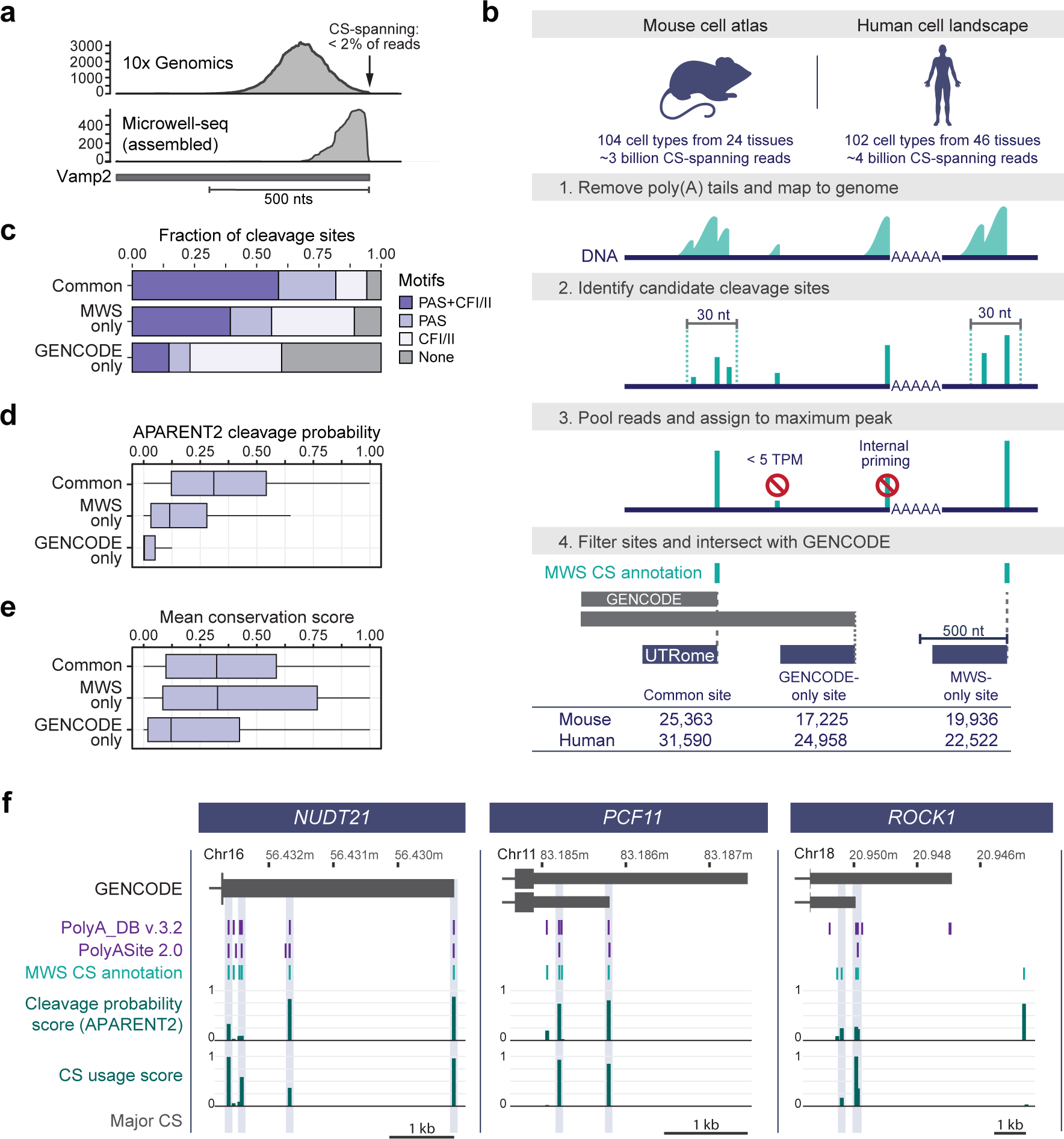
Mapping and characterization of mRNA 3′ end CS in 206 primary cell types. **a,** Read distribution at mRNA 3′ end CS from 10x Genomics compared with MWS data. Shown is the terminal exon of *Vamp2*. nts, nucleotides. **b,** Schematic of CS annotation from MWS data and generation of a truncated UTRome for downstream gene and 3′UTR isoform quantification. **c,** Motif distribution surrounding CS (position 0). PAS (AWTAAA) in [-50,0], CFI binding site (TGTA) in [-100,0], CFII binding site (TKTKTK) in [0,50]. **d,** APARENT2 cleavage probability represents the sum of probabilities in a 30-nt window surrounding the CS. Box shows IQR with median and whiskers 1.5*IQR. **e,** Mean PhastCons score in 30 genomes in a 100-nt window centered on CS, but excluding coding sequences. Box shows IQR with median and whiskers 1.5*IQR. **f,** GENCODE transcript annotations depicting the last exons of *NUDT21*, *PCF11*, and *ROCK1*. Shown are chromosome coordinates (hg38), CS from existing PAS databases and our MWS CS annotation, together with APARENT2 cleavage probability scores and CS usage scores. Major CS are indicated by the grey boxes.

To assess the quality of the novel CS, we analyzed them for binding sites of the CPA machinery. Functional CS typically contain a PAS, an upstream cleavage factor (CF) I binding site and a downstream CFII binding site^1^. We also calculated APARENT2 scores which infer cleavage probability derived from a residual neural network model that was trained on data from a massively-parallel reporter assay^39,40^. Moreover, we analyzed PhastCons DNA sequence conservation surrounding CS^41^. We observed that novel CS have slightly weaker sequence contexts and lower predicted cleavage rates, but they show similar levels of sequence conservation compared to common sites (Fig. 1c-e, Fig. S1d-f).

Next, we compared our new MWS CS annotation with existing databases for individual genes. For most CS, our data agrees with sites annotated in PAS databases, obtained from 3′ end sequencing data^35,36^. However, GENCODE annotations are often incomplete and lack proximal CS (Fig. 1f, S1g). Most of the novel CS were obtained from cell types that are absent in existing CS databases, as illustrated for *ROCK1* (Fig. 1f). Although the *ROCK1* gene is expressed in 90 cell types, the novel distal CS was only detected in lung alveolar stem cells, lung macrophages, and cord blood hematopoietic stem cells (HSCs), thus making it a cell type-restricted site.

Analyzing CS usage across many cell types revealed that not all CS have similar usage rates, but instead can be broadly categorized into major and minor sites. The CS usage score is the fraction of cell types that use a CS divided by the number of cell types that express the gene and are listed in Tables S1 and S2. We observed that nearly half of CS are minor sites as they are used in less than 10% of cell types that express the corresponding gene (Fig. S1h, S1i). Notably, the most distal CS annotated in GENCODE are often minor sites (Fig. 1f, S1g) and when focusing on major CS of all expressed genes, we observed that most mRNAs have only one or two main 3′ ends (Fig. S1j, S1k).

### scUTRquant uses a truncated UTRome for fast and accurate gene and 3′UTR expression quantification

Next, we incorporated our CS atlas into a new computational pipeline for gene and 3′UTR isoform quantification from raw scRNA-seq data. By leveraging our comprehensive reference CS atlas, we circumvent the need for *de novo* peak calling and its reliance on computationally intensive read mapping to a reference genome. Instead, we built the pipeline around the kallisto-bustools^42^ workflow and implemented calibrations for resolving 3′ end isoforms. To reduce isoform assignment ambiguity, we generated a truncated UTRome for pseudoalignment of 3′ end sequencing data, similar to Diag et al. (2018)^43^ (Fig. 1b). The cut-off for the truncation was empirically determined. We observed that more than 95% of 3′ end reads of tested reference genes map within 500 nucleotides (nt) upstream of CS (Fig. S2a). In order to resolve isoform expression originating from close-spaced CS that are less than 500 nt apart, we employed the expectation maximization algorithm implemented in kallisto-bustools. This approach allowed for the proportional assignment of ambiguous reads^44^. As 3′ end sequencing data violate the assumption of uniformly distributed reads used in the implementation^44^, we investigated to what extent an overlap of quantification windows could induce errors in isoform quantification. We performed simulations to determine the error rate as a function of CS distance. We observed that CS within 200 nt of each other cannot be reliably quantified (Fig. S2b). Therefore, we merge their counts and assign them to the distal CS. We call this pipeline scUTRquant, which is available on our GitHub page (https://github.com/Mayrlab/scUTRquant).

To test the consistency of gene counts obtained from the truncated UTRome with standard processing, we compared values obtained from scUTRquant and CellRanger on several 10x Genomics demonstration datasets^45^. We observed nearly perfect correlations, as evidenced by

Spearman’s rank correlation coefficients (ρ) exceeding 0.99 (Fig. S2c) for UMI counts per cell, and 0.93 for UMI counts per gene (Fig. 2a). Moreover, Louvain clustering results based on gene counts were similar (Fig. S2d, S2e). These results demonstrate that gene counts obtained from scUTRquant are consistent with the current gold standard analysis tool.

**Figure 2.**
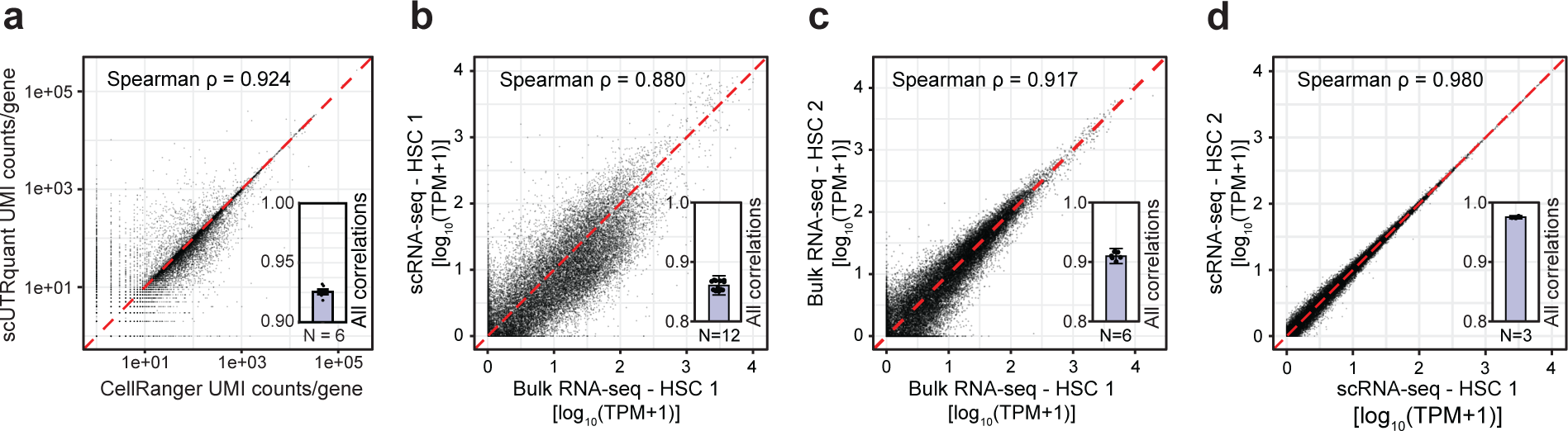
scUTRquant-derived gene and 3′UTR expression is precise and accurate. **a,** Correlation of unique molecular identifiers (UMIs) per gene obtained by scUTRquant and CellRanger for mouse heart 10x Genomics demonstration dataset. Inset: Additional Spearman correlations for six mouse 10x Genomics demonstration datasets. The Spearman’s ρ is shown with mean and SE. **b,** Correlation of 3′UTR isoform counts obtained from FACS-sorted HSCs comparing counts from bulk 3′ end sequencing methods with scUTRquant analysis of scRNA-seq data. Inset: Additional correlations between all pairs of analyzed bulk and scRNA-seq samples. The Spearman’s ρ is shown with mean and SE. **c,** Correlation of 3′UTR isoform counts between two replicates of bulk 3′ end sequencing for FACS-sorted HSCs is shown. Inset: Additional correlations between all pairs of HSC replicates. The Spearman’s ρ is shown with mean and SE. **d,** Correlation of scUTRquant 3′UTR isoform counts between two replicates of scRNA-seq of FACS-sorted HSCs. Inset: Additional correlations between all pairs of HSC replicates. The Spearman’s ρ is shown with mean and SE.

Next, we validated 3′UTR isoform counts by comparing scUTRquant-derived values with 3′UTR isoform counts generated by bulk 3′ end sequencing methods. As only few bulk datasets were generated from well-defined primary cell types, such as embryonic stem cells (ESC) or HSCs, the number of comparisons was limited. For FACS-sorted HSCs, we observed a strong correlation between scUTRquant values and bulk 3′-seq data (Fig. 2b, Spearman’s ρ= 0.88)^46-48^. For ESCs, the correlation was less strong (Fig. S2f, Spearman’s ρ= 0.70), which may be caused by different cultivation conditions^49-52^. Nevertheless, we still consider the level of correlation between the current gold standard method and scUTRquant transcript counts as excellent, considering that the procedures were performed by different laboratories using vastly different methods.

Next, we assessed the reproducibility of 3′UTR isoform counts across biological replicates. Among replicates from scRNA-seq, we observed a substantially stronger correlation than among bulk 3′ end sequencing replicates (Fig. 2c, 2d, Fig. S2g, S2h)^46-52^. Moreover, scRNA-seq-derived biological replicates were highly consistent, even when they were sequenced by different laboratories (Fig. 2d, Fig. S2h)^47,48,51,52^. These results reveal a better accuracy and substantially higher precision, indicating that quantifying 3′UTR isoforms from scRNA-seq data should become the new standard.

### scUTRquant and scUTRboot provide a workflow for 3′UTR analysis from scRNA-seq data

We conceived scUTRquant as part of a broader workflow for analyzing 3′UTR isoforms from scRNA-seq data (Fig. 3a). scUTRquant takes as input either raw scRNA-seq data (in FASTQ format) or output from other pipelines (in BAM format), together with a kallisto index from a truncated CS annotation file, called a UTRome. The comprehensive human and mouse UTRomes are included as default options. However, to provide flexibility for use with other organisms, we developed a Bioconductor package ‘txcutr’^53^ that can generate compatible indexes given any GFF or GTF transcriptome annotation (Online methods). The scUTRquant pipeline provides quality control reports for all samples and outputs gene counts, 3′UTR isoform counts, or both. Count matrices are formatted as Bioconductor ‘SingleCellExperiment’ objects that can be readily used for other scRNA-seq analysis applications. Moreover, previously established cell type annotations can be provided as input and scUTRquant will attach these as column data.

**Figure 3.**
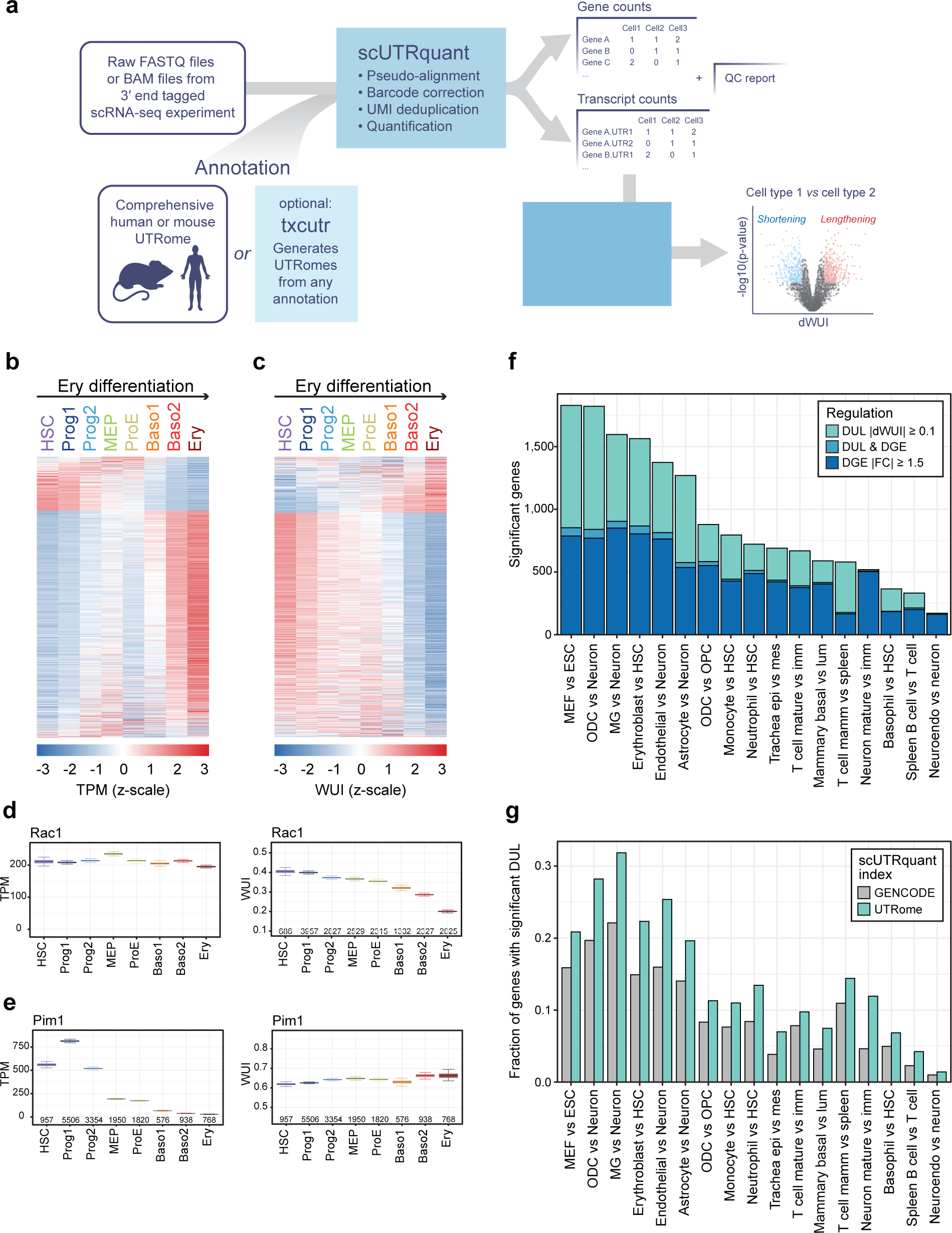
Application of scUTRquant to diverse cell types demonstrates that changes in gene expression and 3′UTR length are independent axes of gene regulation. **a,** Schematic of the scUTRquant pipeline. Inputs are scRNA-seq raw data together with a truncated CS annotation file. The txcutr tool generates truncated UTRomes for any genome. Outputs are quality control parameters and count matrices for gene and 3′UTR expression. The 3′UTR count output can then be used as input for scUTRboot to identify significant differences across known cell types. **b,** Heatmap showing DGE between HSC and Ery. DGE was tested with Welch *t-*test (fold change > 1.5, q-value < 0.05), with 1,059 genes increasing and 252 genes decreasing expression between Ery and HSC. Prog1, progenitor cell type 1, Prog2, progenitor cell type 2, MEP, myeloid-erythroid progenitor, ProE, pro-erythroblast, Baso1, basophilic erythroblast 1, Baso2, basophilic erythroblast 2, Ery, polychromatic erythroblast. **c,** As in **b**, but shown is DUL, calculated with scUTRboot’s WUI bootstrap test (difference in WUI > 0.10, q-value < 0.05). Across all cell types in the differentiation trajectory, 3′UTR lengthening and shortening is observed in 362 and 1,275 genes, respectively. **d,** Example gene (*Rac1*) with significant DUL, but not significant DGE. Bootstrap resampling of cells was used to estimate 95% confidence intervals on the mean TPM (left) and mean WUI (right) for each cell type. Number of cells expressing the gene are indicated along the x-axis. **e,** As in **d**, but example gene (*Pim1*) with significant DGE, but not significant DUL. **f,** As in **b** and **c**, but pair-wise comparisons were performed. Shown is the number of genes with significant DGE and DUL and the genes with both changes for each pair-wise comparison. MEF, mouse embryonic fibroblast; ODC, oligodendrocyte; MG, microglia; OPC, oligodendrocyte precursor; epi, epithelial cell; mes, mesenchymal cell; imm, immature; lum, luminal cell; mamm, mammary; neuroendo, neuroendocrine. **g,** As in **f**, but the fraction of genes with DUL is shown, when using two different CS annotations as input for scUTRquant. The number of DUL was normalized to the number of expressed multi-UTR genes to demonstrate an increase in DUL even after normalizing to the number of tests performed.

For identification of statistically significant changes in 3′UTR isoforms, we implemented a companion R package, called scUTRboot^54^ (Online methods). It provides a flexible set of non-parametric testing procedures to test for changes in APA or IPA, and directional 3′UTR changes (shortening or lengthening).

### Classification of genes into single- and multi-UTR genes

To demonstrate how the scUTRquant pipeline can be used to gain new biological insights, we applied it to scRNA-seq data from 474 cell types in human and mouse. We processed the Tabula Sapiens dataset, which represents 355 unique cell types^55^. Of 16,185 detected protein coding genes, 8,056 expressed a single 3′UTR isoform, while 8,129 genes were classified as multi-UTR genes, corresponding to 50% of human genes (Fig. S3a, Table S3). We classified a gene as multi-UTR when it has at least two CS in the last exon, resulting in 3′UTR isoforms whose relative expression is at least 10% of all 3′UTR counts in at least one cell type. Independently of single- or multi-UTR genes, we identified 4,113 (25%) genes that generate IPA isoforms, thus changing the encoded protein (Fig. S3b, Table S3).

Similarly, we processed scRNA-seq datasets comprising 119 mouse cell types obtained from Tabula Muris, brain, ESCs, and bone marrow^47,48,51,56,57^. Among the 16,195 expressed protein coding genes, we classified 6,766 (42%) as multi-UTR genes and we identified 1,869 (12%) genes that generate IPA isoforms (Fig. S3c, S3d, Table S4). In general, we observed that the majority of genes that are expressed in few cell types only generate one 3′UTR isoform, whereas most genes with broad expression patterns are classified as multi-UTR genes. Interestingly, true ubiquitously expressed genes (detected in more than 88% of cell types) are also more likely classified as single-UTR genes (Fig. S3e, S3f).

### Coordinated 3′UTR length changes during differentiation

Next, we used the scUTRquant pipeline to re-analyze a published bone marrow dataset with the aim of identifying cell-type specific differences in 3′UTR length during red blood cell (Erythroblasts, Ery) differentiation from HSCs^47,48^. To classify 3′UTR changes into shortening or lengthening, we used a weighted UTR index (WUI)^58^. For genes with two or more 3′UTRs, isoform usage is weighted based on order of occurrence, with the shortest and longest isoforms being assigned weights of 0 and 1, respectively (Fig. S3g). The higher the WUI value of a gene, the more of its expression is derived from long 3′UTRs. For example, for genes with two 3′UTRs, the WUI represents the fraction of UMI counts that map to the longest 3′UTR isoform. Along the Ery differentiation trajectory, we identified 1,311 genes with differential gene expression (DGE) using a minimum fold-change of 1.5 as well as 1,637 genes with differential 3′UTR length (DUL), considering WUI changes of at least 0.1 (Fig. 3b-e). Along the Ery differentiation trajectory, both DGE and DUL changes were gradual and coordinated, and they were equally widespread.

### Gene expression and 3′UTR length represent independent axes of gene regulation

To examine if changes in gene expression and 3′UTR length affect the same genes, we focused on the pair-wise comparison between HSC and Ery. We identified 876 DGE and 873 DUL changes (Fig. S3h). Only 103 genes simultaneously changed both gene expression and 3′UTR length, indicating that a gene either changed its expression or it changed its 3′UTR length (Fig. 3f).

To examine the relationship between changes in gene expression and 3′UTR length in other cell types, we determined DGE and DUL in additional differentiation pathways and cell type comparisons. In most pairwise comparisons, ranging from ESC to blood cells and brain cell types, we observed similar numbers of gene expression and 3′UTR length changes (Fig. 3f, S3i). Importantly, the genes that changed their mRNA abundance or their 3′UTR length had little overlap, which ranged from 0-7.4%, considering both types of changes (Fig. 3f). The degree of overlap was consistent with statistical independence in 14/17 sample pairs (Table S5). The overlap between the two groups was still minimal, even when we lowered the cutoff for the expression fold change to 1.25 (Fig. S3i, Table S5). To further exclude a bias caused by data thresholding, we calculated Pearson correlation coefficients across all pairwise comparisons.

The average correlation coefficients were estimated to be less than 0.1, and 3/17 comparisons yielded no significant correlation between the gene expression and 3′UTR length variable (Fig. S3j, Table S5). These data demonstrate that for most genes, expression and 3′UTR length changes represent indeed two independent axes of gene regulation (Table S5).

### The new CS atlas increases detection of differential 3′UTR length by 40%

To investigate whether the new CS annotation can resolve more differential 3′UTR events, we calculated the number of DUL changes when using GENCODE annotations compared with using our UTRome. When using the new CS annotation, we were able to test 20%-80% (median 70%) more multi-UTR genes and detected 110%-220% (median 150%) more differential 3′UTR events (Fig. S3k). When normalizing for the number of expressed multi-UTR genes that were analyzed, we still observed a median 40% higher fraction of genes with differential 3′UTR events compared to using GENCODE annotations only (Fig. 3g). This shows that our more comprehensive CS annotation substantially increases the number of genes with detectable and significant changes in 3′UTR usage across diverse cell types.

### 3′UTR analysis of a Perturb-seq dataset identifies novel APA regulators

We anticipate that the most common application for scUTRquant will be the quantification of 3′UTR isoforms across cell types. To demonstrate additional uses, we applied scUTRquant to a Perturb-seq dataset containing knockdown experiments for 2,134 essential genes to identify known and novel regulators of APA^59^. To validate this approach, we plotted global APA and IPA changes after knockdown of known regulators, including the CPA machinery and the PAF complex. The observed changes recapitulated previously published results and showed that knockdown of core CPA factors causes overall 3′UTR lengthening, whereas knockdown of CFI strongly induces 3′UTR shortening (Fig. S4a-h)^34,60,61^. Knockdown of the PAF complex also increased IPA, which is consistent with its known role as positive regulator of transcription elongation (Fig. S4a-h)^8^. The effect of each perturbation on global 3′UTR shortening or lengthening is reported in Table S6.

To identify novel APA regulators, we calculated a z-scaled difference in WUI (dWUI) between each perturbation and the group of 97 non-targeting controls, followed by clustering on the perturbations and the target genes (Fig. 4a). Our clustering analysis identified 18 perturbation clusters, which contain groups of factors, that when knocked down in K562 cells, caused similar patterns of 3′UTR shortening or lengthening in specific groups of target genes (Fig. 4a, 4b, Table S7). Among the perturbation clusters, we identified several known APA regulators, including members of the CPA machinery as well as splicing, nuclear export, and transcription elongation factors. We also observed several clusters that contain members of large protein complexes, including CCT, nuclear exosome, proteasome, and the ribosome (Fig. 4a, 4b).

**Figure 4.**
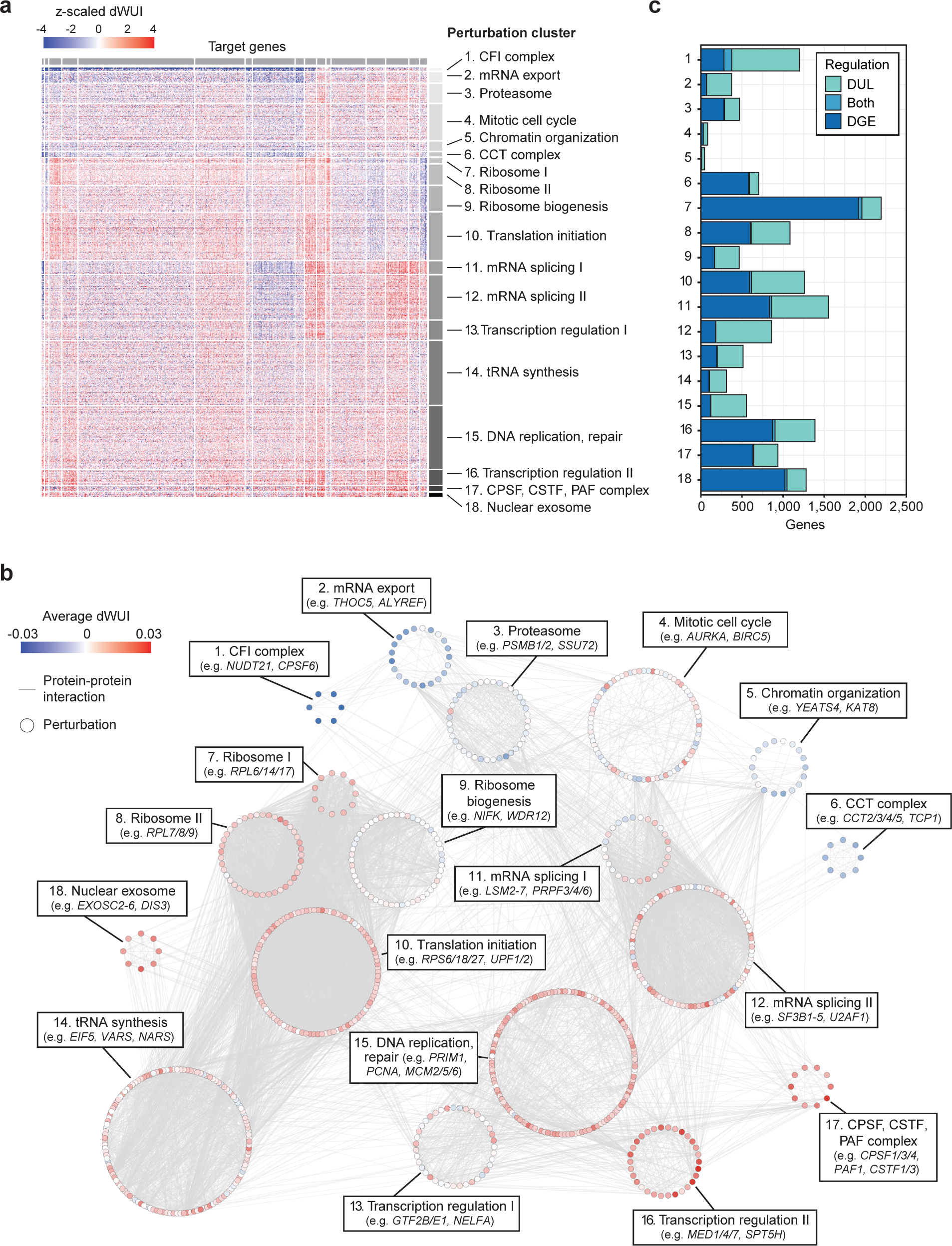
Perturbation clusters that cause similar patterns of 3′UTR changes in specific groups of target genes. **a,** Clustering of essential gene perturbations on z-scaled dWUI values identifies 18 clusters with gene perturbations causing distinct 3′UTR expression patterns. Gene perturbations ordered by perturbation cluster are located on the vertical axis (836 genes) and target genes are presented on the horizontal axis (883 genes). Blue colors indicate shifts towards expression of shorter 3′UTRs upon perturbation, whereas red colors indicate 3′UTR lengthening upon perturbation. **b,** Components of 18 perturbation clusters from **a** are shown in a protein-protein interaction network based on data from the STRING database. Nodes representing individual perturbations were arranged by cluster, and average dWUI values for each perturbation are indicated by node colors. **c,** Bar plot showing the number of genes with DGE and DUL in each perturbation cluster (as in **a**). DGE was determined by two-sided Mann-Whitney test on all genes with a minimum expression of 5 TPM, showing a fold-change > 1.5 with q-value < 0.05. DUL was tested on multi-UTR genes with a minimum expression of 5 TPM and a q-value < 0.05.

As the majority of these factors were previously found to be involved in gene expression regulation^59^, our data suggests that gene and 3′UTR isoform expression are controlled by the same factors (Fig. 4c). To identify whether specific complexes have a bias towards gene or 3′UTR regulation, we intersected DGE and DUL for each perturbation cluster. Whereas perturbing the core CPA machinery mostly affected gene expression, CFI knockdown was biased towards 3′UTR regulation (Fig. 4c). Again, we found that across all multi-UTR genes and perturbation clusters, the correlation between UTR change and gene expression was weak (average *R* ≈ −0.06, Fig. S4i).

### Reduction of ribosomal proteins causes widespread APA changes

To obtain deeper insights into APA regulation, we separated the changes induced by each perturbation cluster into 3′UTR shortening or lengthening (Fig. 5a). The strongest APA regulator was CFI, whose knockdown nearly exclusively induced 3′UTR shortening, as was reported previously for other contexts^34,60,61^. Perturbation of mRNA export factors also predominantly induced 3′UTR shortening (Fig. 5a), indicating that presence of CFI or export factors promote full length mRNA isoform expression. In addition to perturbing CFI, knockdown of splicing factors caused the largest numbers of APA changes (Fig. 5a). This was followed by perturbation clusters that contain transcription factors, translation initiation factors, and the ribosome (Fig. 5a). Although inhibition of translation is known to be associated with mRNA decay^62^, it was surprising to find that inhibition of factors that target the mRNA region common to both 3′UTR isoforms can differentially affect their expression.

**Figure 5.**
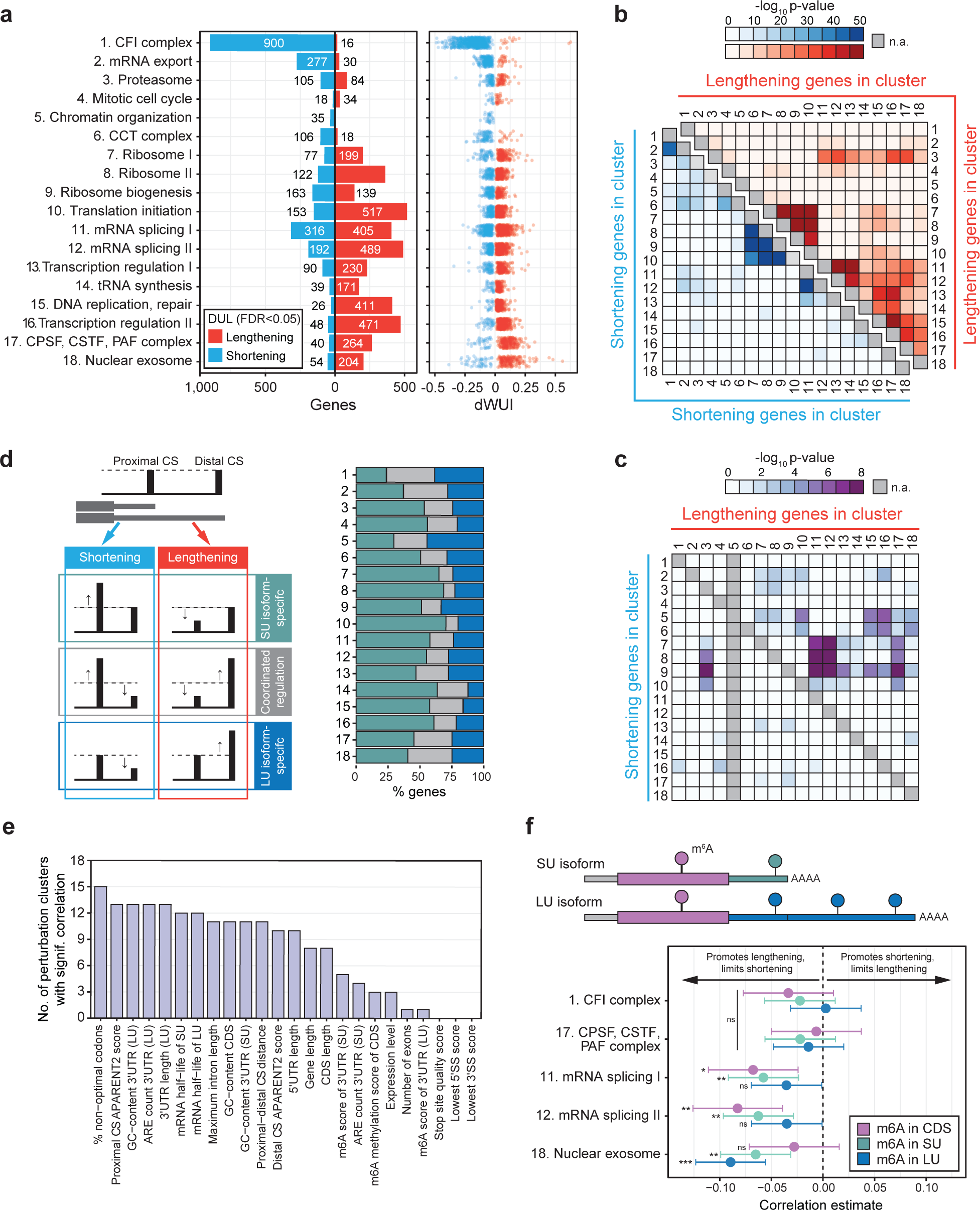
Regulatory logic of 3′UTR changes in 18 perturbation clusters. **a,** Left panel: Bar plot showing the number of genes with significant 3′UTR shortening (blue) or lengthening (red) in each perturbation cluster relative to 97 samples expressing non-targeting guide RNAs. DUL was performed using a two-sided Mann-Whitney test with a q-value < 0.05. Right panel: Dot plot showing the average WUI difference for significant genes in the cluster. **b,** Heatmap depicting the probability of gene overlaps observed between perturbation clusters with synergistic regulation for 3′UTR lengthening (upper right, red) or 3′UTR shortening (lower left, blue), compared to what would be expected by chance, calculated using Fisher’s exact test. **c,** As in **b**, but for antagonistic regulation of gene sets. **d,** Left panel: Schematic diagram that illustrates the potential mechanisms leading to 3′UTR shortening and lengthening. In both cases, regulation can occur in a transcript-specific manner, affecting only one of the isoforms, or in a compensatory fashion where both isoforms change in a coordinated manner. Right panel: The mechanism of change, categorized as either short UTR (SU)-specific (green), coordinated (grey), or long UTR (LU)-specific (blue) is shown for the significant genes from **a.** Only genes that generate two 3′UTR isoforms were analyzed. Coordinated change was defined as a relative shift of expression of at least 50% from one to the other isoform. **e,** Gene and mRNA features that are significantly associated with 3′UTR changes (dWUI) observed in each perturbation cluster. The number of significant correlations with q-value < 0.05 for each feature are shown. ARE, AU-rich element; CDS, coding sequence; SS, splice site. See also Figure S5c and Table S8. **f,** Pearson correlation coefficients between m6A methylation marks of all multi-UTR genes with TPM >5 and a subset of perturbation clusters are shown for different mRNA regions. Error bars show 95% CI. Significance of correlations are indicated with ns (not significant) if q-value > 0.05, ** q-value < 0.01, *** q-value < 0.001. SU (short 3′UTR) isoform, LU (long 3′UTR) isoform.

Each perturbation cluster regulated distinct sets of target genes. Next, we characterized the extent of overlap in target genes that are regulated in the same or opposite direction across perturbation clusters (Fig. 5b). As expected, perturbation clusters that contain genes with related molecular functions showed the strongest overlap. For example, all four clusters with genes related to ribosome function induced 3′UTR shortening (or lengthening) of common target gene sets. The two splicing clusters also affected similar genes. Interestingly, we observed that the targets of CFI and mRNA export factors were highly overlapping (Fig. 5b), which suggests that these factors may be part of a common pathway. Moreover, whereas perturbation of splicing, transcription, and CPA factors induced 3′UTR lengthening of similar genes (Fig. 5b), perturbation of the ribosome caused 3′UTR shortening of this gene set (Fig. 5c).

### Most APA changes affect the shorter 3′UTR isoform

APA changes are commonly described as 3′UTR shortening or lengthening, but these patterns arise through various types of isoform changes (Fig. 5d). 3′UTR shortening can occur through exclusive upregulation of the shorter isoform, exclusive downregulation of the longer isoform or through a combination of the two events (Fig. 5d). Across all changes that affect genes with two 3′UTR isoforms, we observed that compensatory changes, where one isoform increases at the expense of the other isoform are not the dominant mode of regulation (Fig. 5d). Most often, the shorter 3′UTR isoform changes abundance (Fig. 5d). Interestingly, genes that were affected by many perturbation clusters had significantly weaker proximal CS (Fig. S5a-c). Taken together, these results suggest that most regulatory events happen at the proximal CS.

### Gene features of the coding sequence frequently affect expression of alternative 3′UTRs

To better understand the logic why specific groups of genes are affected by particular perturbation clusters, we identified gene and mRNA features that correlate with the extent of 3′UTR changes, stratified by each perturbation cluster (Fig. 5e, Fig. S5d). For example, a high percentage of sub-optimal codons promoted 3′UTR shortening in most perturbations (Fig. 5e, Fig. S5d). In addition to codon optimality, we identified several additional features that are intrinsic to the gene or mRNA region common to both 3′UTR isoforms. These features include maximum intron length and coding region length (Fig. 5e, Fig. S5d, Table S8).

Another feature that correlated with APA changes was m6A methylation. We observed that the presence of the m6A methylation mark in the mRNA mainly supports long 3′UTR isoform expression, as was reported previously^63,64^. Interestingly, our data revealed that the relative position of the modification mark determines the primary mechanism that causes 3′UTR lengthening. For example, 3′UTR isoform expression changes caused by perturbations in splicing factors was significantly correlated with the presence of m6A modifications in the coding region or the short 3′UTR. On the other hand, m6A present in the long 3′UTR influenced differential transcript expression after exosome knockdown (Fig. 5f, Fig. S5d).

## Discussion

Our study presents the first comprehensive annotation of human and mouse mRNA 3′ end CS using data from hundreds of primary cell types^37,38^. More than 200 cell types derived from all major organs and developmental stages were included in the analysis. We performed rigorous quality control and filtering for internal priming and show that the novel CS are of high quality. We extend current GENCODE CS annotations by 40%. In our new annotation, called MWS CS annotation, we integrated the novel CS with GENCODE annotations and provide the most comprehensive CS annotation for human and mouse to date.

In addition to expanding CS annotations, our large-scale analysis on cell type-specific CS usage allowed us to categorize CS into major and minor sites. This revealed that most genes only have one or two major CS. As their locations within the mRNA are identical across cell types, our data suggests that CS locations are intrinsic gene features. In our opinion, the locations of major CS should be used to re-define 3′UTR boundaries in widely used transcriptome annotations. We observed that the GENCODE and Refseq databases have been expanding their 3′UTR annotations, which has led to the presence of very long 3′UTRs for many genes in human and mouse (Fig. 1f, S1g). Although these CS may be used in a few rare cell types, they represent minor CS that should not be used for the definition of 3′UTR length under most conditions. Our categorized CS annotation clarifies transcriptome-wide mRNA 3′ end boundaries and increases our understanding of cell type-specific differences in mRNA 3′ ends.

In addition to the CS annotation, we developed an open-source computational pipeline, that makes it simple to quantify gene and 3′UTR isoform expression from new or existing scRNA-seq. By building on the kallisto-bustools workflow, we provide a fast and direct means of quantification that is compatible with most 3′-end tag scRNA-seq libraries. We established optimal parameters for reliable quantification and implemented these as default settings. With our pre-defined CS annotation, scUTRquant quantifies a consistent set of 3′UTR isoforms, making it easier to integrate datasets. Coupled with scUTRboot, significant differences in 3′UTRs across samples are identified, which allows the integration of 3′UTR quantification into standard scRNA-seq data analysis.

To demonstrate how scUTRquant can be used to gain new biological insights, we analyzed the global 3′UTR changes in a recent Perturb-seq data set that contains over 2,000 knockdown experiments^59^. We established a comprehensive catalog of 3′UTR regulators that validated known factors and significantly expanded our knowledge on the mechanisms of APA regulation. For instance, in addition to factors of the CPA machinery, we observed that splicing has a substantial and widespread impact on expression of alternative 3′UTRs. Moreover, we identified the ribosome and translation initiation as major influencers of differential 3′UTR expression.

While translation has long been recognized as an important regulator of mRNA decay^62^, we provide evidence for translation-dependent differential turnover of mRNAs with alternative 3′UTRs. This may be surprising as the coding regions of alternative 3′UTR isoforms are identical. Our data suggest that 3′UTR-bound RNA-binding proteins communicate with the translation environment to trigger translation-dependent mRNA decay. This may allow the ribosome to integrate signals from different parts of the mRNA, including codon usage from the coding sequence and 3′UTR-bound RNA-binding proteins.

We further applied the scUTRquant pipeline to 3′UTR analysis across 474 human and mouse cell types. By integrating changes in gene expression and in 3′UTR length across many cell types, our analysis revealed that mRNA abundance and in 3′UTR length are two independent measures of gene output (Fig. 3f). Our results confirm previous observations that were obtained from a limited number of cell types^2,23-27^. Here, we demonstrate that only approximately 10% of differential 3′UTR events are associated with changes in gene expression, which indicates, that in most cases, APA is not a mechanism for gene expression regulation. Importantly, we revealed that only approximately half of gene output changes mRNA abundance, whereas the other half changes 3′UTR length, and therefore controls the presence or absence of regulatory motifs in mRNAs.

In addition to regulating abundance, 3′UTRs are known as major regulators of mRNA localization^22^, even in non-neuronal cell types such as enterocytes, endothelial cells, mammary epithelial cells, and HEK293T cells^14,20,65-67^. Moreover, 3′UTRs have emerged as important regulators of mRNA-dependent assembly of proteins complexes^12,13,17,18,21^. Taken together, our large-scale analysis revealed that about half of gene regulatory events go undetected when using standard gene expression analysis pipelines. Rather than controlling the abundance of transcripts, these changes may instead predominantly modulate the spatial organization of the transcriptome.

## Supporting information

Table S1 - Human MWS CS Annotations

Table S2 - Mouse MWS CS Annotations

Table S3 - Human Gene Annotations

Table S4 - Mouse Gene Annotations

Table S5 - DGE versus DUL

Table S6 - Global WUI and IPA Changes

Table S7 - APA Perturbation Clusters

Table S8 - Cluster-Feature Correlations

## Acknowledgements

This work was funded by NIH training grant T32GM083937 to M.F., the NIH Director′s Pioneer Award (DP1-GM123454), an R35GM144046 NIH grant, a grant from the Pershing Square Foundation, William Ackman, and Neri Oxman, and the MSK Core Grant (P30 CA008748) to C.M. We thank Quaid Morris and all members of the Mayr lab for helpful discussions. We also thank Andrew Grimson, Lucy Skrabanek, and Steve Lianoglou for useful comments on the manuscript.

## Author contributions

M.M.F. and C.M. conceived the project. M.M.F. developed and implemented all new computational tools and processed all datasets. S.M. analyzed the Perturb-seq data. All authors designed the experiments and wrote the manuscript.

## Declaration of Interests

The authors declare no competing interests.

## List of Supplemental Tables

**Table S1. MWS CS annotations and CS classification of the human transcriptome.**

**Table S2. MWS CS annotations and CS classification of the mouse transcriptome.**

**Table S3. Single-, multi-UTR, IPA genes in human.** Reported is also 3′UTR length of each isoform.

**Table S4. Single-, multi-UTR, IPA genes in mouse.** Reported is also 3′UTR length of each isoform.

**Table S5. Changes in gene expression and 3′UTR length are independent parameters.**

**Table S6. Global 3′UTR and IPA changes in each perturbation.**

**Table S7. APA regulators contained in each perturbation cluster.**

**Table S8. Gene and mRNA features that correlate with 3′UTR changes in each perturbation cluster.** Reported is also a comprehensive list of external data sources that were used for the analysis.

## List of available data files

**GTF and BED files for the MWS CS annotation in human and mouse** are available at https://doi.org/10.6084/m9.figshare.23549526.

## Online methods

### Cell type-specific identification of mRNA 3′ ends from primary cells using MWS data

<u>CS identification.</u> FASTQ files of MWS data from the Mouse Cell Atlas v1.1^37^ and the Human Cell Landscape^38^ were downloaded and then assembled using PEAR v0.9.6 with settings ‘-n 75 -p 0.0001’. Cell and UMI barcodes were extracted from assembled reads and placed into read headers using umi_tools v1.1.2; remaining poly-T regions at the 5′ end of assembled reads were trimmed using cutadapt v3.5 with arguments ‘--front=’T{100}’ -g=’T{12}’ -n 10 -e 0’, retaining only sequences with minimum length of 21 nts. Reads were aligned with HISAT v2.2.1 to the mm10 and hg38 genomes, respectively. Cell type annotations were used to demultiplex sample-level BAMs to cell-type-level for each dataset. Per cell-type strand-specific coverage at the 5′ ends of aligned reads was computed using the ‘genomecov -dz -5’ command of BEDTools v2.30. Per cell-type, all entries within 30 nt radius were merged to the local mode and retained when > 5 reads per million. Cell type coverages per strand were subsequently summed with GNU′s datamash v1.7, and then a final pass of merging to the local mode within a radius of 30 nt was performed to harmonize minor variations in cell-type-level CS identification. The resulting sites were considered as CS candidates (mouse: *N* = 170,617; human: *N* = 150,191).

<u>CS filtering.</u> Candidate CS were intersected with 40 nt intervals centered at 3′ ends of GENCODE vM25 and v39 transcripts with positively identified 3′ ends (excluding tag ‘mRNA_end_NF’), respectively. Intersecting sites (mouse: *N* = 27,872; human: *N* = 31,026) were classified as “validated”; non-intersecting sites were subsequently intersected with 40 nt intervals centered at cluster centers in the PolyASite v2.0 Mus musculus and Homo sapiens atlases using all clusters surpassing a 3 TPM threshold^36^. Intersecting sites (mouse: *N* = 30,020; human: *N* = 29,897) were classified as “supported”; non-intersecting sites were filtered through cleanUpdTSeq v1.32 with maximum posterior probability of 0.0001 of being an internal priming site^68^. Passing sites (mouse: *N* = 22,824; human: *N* = 18,055) were classified as “likely”. The union of “supported” and “likely” CS was formed and each site was annotated according to the GENCODE vM25 and v39 annotations, with one of the ordered labels: “three_prime_UTR”, “five_prime_UTR”, “exon”, “intron”, “extended_five_prime_UTR”, “extended_three_prime_UTR”, or “intergenic”, where the existing 5′ ends of transcripts were extended 1 kb upstream and existing 3′ ends of transcripts were extended 5 kb downstream.

<u>Generation of the MWS CS annotation and transcriptome truncation.</u> The GENCODE vM25 and v39 annotations were filtered for protein-coding transcripts with known 3′ ends. CS with a “three_prime_UTR” label were intersected with these transcripts and new transcript versions ending at the CS were generated (“upstream”). All protein-coding transcripts with known 3′ ends were extended by 5 kb downstream, intersected with the “extended_three_prime_UTR” set of CS, and new transcript versions ending at the CS were generated (“downstream”). All transcripts (GENCODE, upstream, downstream) were truncated to include 500 nts from their 3′ end. Truncated transcripts with fewer than 50 nts difference were reduced to a single representative copy, with prioritization for downstream sites. The collection of remaining truncated transcripts was exported to GTF and the corresponding sequences to FASTA. This expanded the GENCODE annotations with 19,936 and 22,522 additional 3′UTR isoforms, respectively. The new annotation is called MWS CS annotation. The GTF and BED files are provided as a figshare link (https://figshare.com/s/0709e2551cc1ee4c4941).

### Characterization of CS in the MWS CS annotation

<u>Sequence motifs surrounding CS.</u> DNA sequence in a window of 1,000 nt centered at each CS was extracted. For each motif, the center positions of all occurrences were determined across all sequences. Smoothed density is computed with ggplot’s ‘stat_density’, with global mode scaled to 1.

<u>APARENT2 cleavage probabilities.</u> DNA sequence in a window of 205 nt centered at each CS in the MWS CS annotation was extracted and used as input to APARENT-ResNet v1.0.2^40^. Cleavage probability for a site was computed as the sum of probabilities output by APARENT-ResNet in the 30 nt window centered at the CS.

<u>DNA sequence conservation scores.</u> Conservation scores were extracted in 100 nt windows centered at CS, with annotated coding sequence regions excluded, using the Bioconductor package ‘GenomicScores’ v2.10.0 with databases “phastCons30way.UCSC.hg38” and “phastCons60way.UCSC.mm10”. Means were computed for each set of scores from the window.

<u>Comparisons with existing CS databases.</u> Plots in Figures 1f and S1g included only clusters from PolyASite 2.0 with a minimum TPM of 1 or relative usage rate above 0.05. PolyA_DB v3.2 CS were similarly filtered but using a 1 RPM threshold. GENCODE (human v39; mouse vM25) were filtered to exclude transcripts denoted with “mRNA_end_NF”.

<u>The CS usage score classifies CS into major and minor sites.</u> The CS usage score is the fraction of cell types that use a CS in the data from MWS divided by the number of cell types that express the gene. If a CS is used in < 10 cell types where the gene is expressed, it is considered a minor CS. All CS together with their CS usage scores are reported in Tables S1 and S2 (for human and mouse CS annotations).

### Definition of scUTRquant parameters

<u>Empirical distributions of 10x Genomics peak width.</u> A set of 56 peaks located at the 3′ ends of transcripts was manually curated by examining the genomic alignments of 10x Genomics Chromium v2 samples from the Tabula Muris dataset^56^. Peaks were selected for absence of splice sites, potential internal priming sites (A-rich regions), and nearby alternative CS in the immediate 800 nts upstream of the annotated CS. The coverage of 5′ ends of reads was extracted with the ‘bedtools genomecov -5’ command for each sample from the Tabula Muris dataset and the distance from the annotated CS of the corresponding transcript was computed. For each sample, the 95^th^ percentile for distance from the 3′ end across all genes was computed. Additionally, for each gene, the 95^th^ percentile for distance from the 3′ end across all samples was computed (Fig. S2a). Analysis code available at https://github.com/Mayrlab/tmuris-peaks.

<u>Kallisto transcript quantification resolution.</u> To resolve CS nearer than 500 nt apart we enabled the expectation maximization algorithm implemented in kallisto-bustools to proportionally assign ambiguous reads^44^. As 3′ end sequencing data violate the assumption of uniformly distributed reads used in the implementation^44^, we investigated to what extent overlapping isoforms might induce quantification errors. We performed simulations to determine the error-rate as a function of CS distance.

The sequence of the Ensembl transcript Rac1-201 (ENSMUST00000080537) was used as the basis for a two-isoform transcript expression simulation. The first simulated isoform (“distal”) used the annotated 3′ end; the second (“proximal”) was created by removing specified intervals from the 3′ end. For each round of simulation, samples of read distances from the 3′ end of each transcript were generated according to a discretized gamma distribution with mean 300 and standard deviation of 100. Reads of 100 nts were generated using the respective transcript sequences and the randomly sampled positions. The ‘kallisto quant’ command was used to estimate transcript abundance, using the parameters ‘--single -l1 -s1 --fr-stranded -- pseudobam’ and truncated versions of the transcripts as index. Relative error for each transcript was computed using estimated and true abundances. A parameter sweep was performed with all combinations of the following parameters: (a) CS distances between [50-700] with 50 nt steps; (b) truncated transcript lengths [350-600] with 50 nt steps; (c) proximal counts {50,100}; (d) distal counts {50,100}. Each parameter combination was simulated for 10 replicates. Final resolution was selected based on mean relative errors approaching zero. We concluded that CS within 200 nt of each other cannot be reliably discriminated when quantifying (Fig. S2b), therefore, we configured the scUTRquant pipeline to merge their counts and assign them to the distal CS. Analysis code available at https://github.com/Mayrlab/kallisto-overlap.

<u>Kallisto customization and scUTRquant settings.</u> The ‘kallisto bus’ command of kallisto version

0.46.2 was extended to support strand-specific pseudoalignment for both FASTQ and BAM input files (https://github.com/mfansler/kallisto/releases/tag/v0.46.2sq). All 10x Genomics 3′ end datasets were pseudoaligned with ‘kallisto bus --fr-stranded’. Cell barcodes for the corresponding technology version (v2 or v3) were used as whitelists for the ‘bustools correct’ command. Truncated isoforms in the same gene with 3′ ends nearer than 200 nts apart (mouse: *N* = 18,057; human: *N* = 29,574) were merged in the ‘bustools count’ step.

### txcutr generates truncated UTRomes for any genome annotation as input for scUTRquant

The default CS annotation for the scUTRquant pipeline is the human or mouse UTRome which contains the MWS CS annotation. Additional truncated UTRomes for any genome annotation can be created with the Bioconductor ‘txcutr’ package and used as CS annotation input for scUTRquant. The Bioconductor ‘txcutr’ package generates truncated GTF annotations, FASTA sequences, and merges tables^53^.

To investigate how many new differential 3′UTR events can be detected when using the new mouse MWS CS annotation compared with GENCODE vM25, we used ‘txcutr’ v0.99.0 (equivalent to Bioconductor version 1.0.0). This index was generated with a 500 nt truncation length and a merge distance of 200 nts. In brief, the GENCODE vM25 annotation was first pre-filtered with an AWK script to remove any entries with the ‘mRNA_end_NF’ tag (indicating unvalidated 3′ ends) and restricted to protein-coding transcripts. The txcutr method ‘truncateTxomè clipped all transcripts longer than the specified length, anchored at the 3′ end, intersected the truncated transcripts with the child exons of that transcript, and then redefined the genomic range of the gene to the union of all child transcripts. Transcripts that were identical after truncation were deduplicated to retain only one representative copy, which was annotated with the transcript ID of the transcript with lexicographical priority. The resulting TxDb object was then exported as a GTF and a FASTA file using txcutr’s ‘exportGTF’ and ‘exportFASTÀ methods. Finally, a merge table was generated with txcutr’s ‘generateMergeTablè by further truncating transcripts to the specified merge distance, anchored at the 3′ end, intersecting within the parent gene, and recording the most downstream transcript with which each intersects. Additional specification and implementation details are found in the txcutr documentation. The generation of this index is reproducible from the Snakemake pipeline available at https://github.com/Mayrlab/txcutr-db.

### Validation of scUTRquant-derived gene and 3′UTR isoform counts

<u>CellRanger and scUTRquant UMI count correlations.</u> To test the accuracy of gene counts obtained from the truncated UTRome, we compared gene expression values calculated with scUTRquant and CellRanger on six 10x Genomics 3′ end mouse demonstration datasets, available as FASTQ files from the 10x Genomics website (‘heart_1k_v2’, ‘heart_1k_v3’, ‘heart_10k_v3’, ‘neuron_1k_v2’, ‘neuron_1k_v3’, and ‘neuron_10k_v3’). They were processed through the scUTRquant pipeline using ‘utrome_mm10_v2’ target with default settings. The corresponding filtered HDF5 UMI counts from CellRanger 3.0.0 were also downloaded and loaded as SingleCellExperiment objects in R^45^. For each dataset, only cells (or genes) present in both the CellRanger and scUTRquant results were plotted and used to compute Spearman correlations.

Similarly, three 10x Genomics 3′ end human demonstration datasets (‘pbmc_1k_v2’, ‘pbmc_1k_v3’, and ‘pbmc_10k_v3’) were processed using scUTRquant with the Ensembl Release 93 annotation preprocessed according to the CellRanger 3.0.0 pipeline and truncated to 500 nts using txcutr. Comparisons were performed against CellRanger UMI counts in the same manner as above.

The scripts and input files needed to download and run these datasets were incorporated into the scUTRquant pipeline as examples that users can run following the pipeline documentation.

<u>CellRanger and scUTRquant clustering comparisons using gene counts.</u> For each 10x Genomics dataset, the CellRanger and scUTRquant counts were filtered to common cells. Clustering was performed following Amezquita et al., (2019)^69^. In brief, size factors were computed with the ‘computeSumFactors’ from the ‘scran’ Bioconductor package^70^, and then used to compute normalized log counts. The top 1000 high-variance genes were used to compute the first 20 principal components. Louvain clustering was performed on the cells in this reduced representation. The Adjusted RAND Index (ARI) between the CellRanger and scUTRquant clusters was computed using the ‘aricodè R package^71^.

<u>Comparison of 3′UTR isoform counts obtained by scUTRquant with bulk 3′ end sequencing</u> <u>methods.</u> For scRNA-seq data, we used mouse datasets from FACS-sorted HSCs (clusters 0 and 1) according to cell type annotations from Wolf et al., (2019)^47,48^ and from ESCs^51,52^. The UMI counts were summarized to TPM per sample by aggregating counts across all cells in each sample and normalizing to UMIs per million. Bulk 3′-seq datasets on the same cell types were obtained. For FACS-sorted HSC samples^46^, 3′UTR isoforms were quantified by pseudoalignment of read 2 using the UTRome annotation and ‘kallisto quant’. For bulk ESC datasets^49,50^, TPM values and CS locations were obtained from the PolyASite v2.0 database^36^ and intersected with the UTRome annotation using a 50 nt interval around cluster centers. TPM values between the samples were compared and Spearman’s correlation coefficients were calculated.

### Application of scUTRquant to scRNA-seq data from 474 human and mouse cell types

<u>Classification of single- and multi-UTR genes from 119 mouse cell types.</u> Samples from Tabula Muris, ESC, bone marrow, and brain datasets^47,48,51,56,57^ were quantified for 3′UTR isoform usage following the default settings of scUTRquant with the ‘utrome_mm10_v2’ target and cells were annotated with published cell type annotations. Cell type labels for bone marrow cell types were obtained by combining publicly available transcriptome and proteome information for erythroblast differentiation^48,72^. Cells not previously annotated in published analyses were excluded.

All datasets were merged into one SingleCellExperiment object and counts were size-factor normalized using the ‘computeSumFactors’ method from Bioconductor package ‘scran’^70^. UMI counts were aggregated by cell type and the percentage of isoform usage per gene was computed, excluding isoforms whose 3′ ends were located within a GENCODE-annotated intron of the corresponding gene. For each gene, the number of isoforms with at least 10% usage in at least one cell type were counted. Genes with two or more such isoforms were classified as multi-UTR genes; otherwise, they were classified as single-UTR genes. To identify genes that generate intronic polyadenylation (IPA) isoforms, all mRNA 3′ ends of a transcription unit were included. IPA isoforms were counted if they contained at least 10% of reads of a gene in at least one cell type. The data on mouse single-, multi-UTR, and IPA genes together with the 3′UTR length is reported in Table S4. The Snakemake pipeline for classification is available at https://github.com/Mayrlab/atlas-mm.

<u>Classification of single- and multi-UTR genes from 355 human cell types.</u> 10x Genomics samples from Tabula Sapiens^55^ were downloaded in BAM format and processed similarly to the mouse data, but using the scUTRquant ‘utrome_hg38_v1’ target. The data on human single, multi-UTR, and IPA genes together with the 3′UTR length of each isoform is reported in Table S3. The Snakemake pipeline for classification is available at https://github.com/Mayrlab/atlas-hs.

### Identification of differential 3′UTR isoform usage between samples using scUTRboot

To identify statistically significant changes among 3′UTR isoforms across samples, we developed a companion R package, called ‘scUTRboot’^54^. It provides a flexible set of non-parametric testing procedures to test for changes in 3′UTR isoform or IPA isoform usage, directional changes (3′UTR shortening/lengthening) or focus on usage changes in specific classes of isoforms.

<u>Two-sample bootstrap test with scUTRboot.</u> scutrboot implements two-sample hypothesis testing with a bootstrap strategy for estimating p-values. The ‘twoSampleTest’ function implements three general modes of tests based on the statistic computed across the samples: a Usage Index (UI), a Weighted Usage Index (WUI), and a Wasserstein Distance (WD), also called the Earth Mover′s Distance.

<u>For the UI statistic</u>, users provide an indicator vector (‘featureIndex’), indicating a feature such as short 3′UTR isoform (SU), long 3′UTR isoform (LU), or IPA isoform for each gene. The UI statistic per gene is computed as the difference in the fraction of usage of this isoform in the gene across the two sets of cells. This characterizes the difference across sets of cells for a single feature.

<u>The WUI statistic</u> generalizes the UI statistic for genes with several isoforms by using a weighted mean usage across isoforms. scUTRboot supports the usage of arbitrary weights. Throughout this work, we use a particular form we call the ‘Weighted UTR Index’, where the weights correspond to the positional rank of the CS from 5′ to 3′ scaled to the unit interval. That is, the shortest and longest isoforms are assigned weights of 0 and 1. An example of this is shown in Fig. S3g. Concretely, a two-isoform gene will have weights {0,1}, a three-isoform {0, 1/2, 1}, a four-isoform {0, 1/3, 2/3, 1}, and so forth.

<u>The WUI statistic</u> generalizes the UI statistic for genes with several isoforms, as the isoform usages are weighted based on the order of occurrence from 5′ to 3′. Alternatively, users interested in statistical tests of average 3′UTR length could input the length of each isoform as the weight (not reported).

<u>The WD statistic</u> per gene is computed as half the total difference in all isoform usages in the gene across the two sets of cells. When a gene has exactly two isoforms, the UI and WD statistics are identical in magnitude. For genes with several isoforms, the WD statistic incorporates changes in any isoform.

For each of these modes, p-values per gene are estimated using bootstrap resampling under the null hypothesis that the two sets of cells were sampled from identically distributed populations. Specifically, the union of the two sets of cells is used to sample with replacement sets of cells of the same size as the original samples. For each bootstrap sample, the statistic of interest is computed per gene and the p-value is estimated as the fraction of bootstrap statistics as extreme or greater than the observed statistic, with a pseudocount of 1 included to provide a conservative upper bound for rare events. All tests are two-sided.

scUTRboot includes a ‘minCellsPerGenè option to exclude genes that are not sufficiently coexpressed in the samples to compare with confidence. When this is set, bootstrap samples that do not satisfy this minimum are discarded and the p-value will only be computed from the retained samples. The number of retained bootstraps samples used to estimate the p-value is included in the test results.

<u>Bootstrap mean TPM and WUI estimates.</u> For each cell type, 2,000 bootstrap samples were generated by resampling with replacement from the pool of all cells with that cell type. For each bootstrap sample, two statistics were computed: a TPM value and WUI. The TPM value was computed by averaging the TPM value from across all cells in the sample, by gene.

The WUI value was computed by first summing the transcript counts across cells to pseudobulk and then computing the WUI per gene. Percentile statistics were then calculated for these values across the bootstrap samples to determine the confidence interval on the mean TPMs and mean WUIs

<u>Pairwise two-sample bootstrap tests on the differentiation trajectory from HSC to Ery.</u> scUTRboot was used to perform two-sample WUI tests for all non-IPA isoforms detected in at least one cell type in the differentiation trajectory^47,48^. Tests were performed on all pairs of cell types (8 cell types, 28 unique pairs) using 10,000 bootstrap samples on all co-expressed genes (minimum 50 cells expressing each gene) and corrected for multiple testing using Benjamini-Hochberg procedure. Genes were classified as significant if |dWUI| > 0.10 and q-value < 0.05.

<u>Comparing differential gene expression with differential 3′UTR isoform usage.</u> Differential gene expression was performed on pairs of cell types following Amezquita et al., (2019)^69^. In brief, gene-level UMI counts were log-normalized using size factors and a pseudocount of 1. Differential expression was tested with a Welch *t*-test^73^. All p-values were corrected using the Benjamini-Hochberg procedure and genes were classified as significant if fold-changes exceeded 1.5 in either direction and q-value < 0.05.

To identify genes with differential 3′UTR isoform usage, two-sample WUI tests were performed on all cell type pairs (Fig. 3f) using scUTRboot on size-factor normalized UMI counts. All p-values were corrected using the Benjamini-Hochberg procedure and genes were classified as significant if |dWUI| > 0.10 and q-value < 0.05.

To test if gene expression and 3′UTR isoform usage are independent, for each comparison, all coexpressed multi-UTR genes were classified as either non-significant, DGE only, DUL only, or both. A Chi-Square test for independence was performed on the resulting tabulation.

### Application of scUTRquant to a scRNA-seq dataset with 2,134 perturbations

<u>Processing of the Perturb-seq data set</u>. BAM files for the K562 6-day essential gene Perturb-seq experiment^59^ were processed with scUTRquant using ‘utrome_hg38_v1’ index configured for 10x Chromium 3′ end v3. Cell annotations were extracted from the deposited H5AD object (“K562_essential_raw_singlecell_01.h5ad”) and provided to scUTRquant, which transferred the perturbation annotations required for downstream analysis (https://doi.org/10.25452/figshare.plus.20029387.v1). Cells lacking a published annotation were omitted from further analysis. Cells were summarized to pseudobulk by aggregating counts from cells with identical perturbations. We identified all isoforms in terminal exons with at least 10% mean usage within the 97 non-targeting perturbations, and classified genes with at least two such isoforms as multi-UTR genes in this cell line. This procedure yielded 4,780 multi-UTR genes comprised of 11,129 isoforms in active use. The two highest usage isoforms in each gene were designated short UTR (SU) and long UTR (LU) with respect to their 5′-3′ order. TPM values were calculated for all genes and all SU and LU isoforms. WUI values were computed for each multi-UTR gene in each perturbation. The rate of IPA isoform usage was computed for all genes with expressed IPA isoforms for each perturbation.

<u>Calculation of average dWUI and dIPA values.</u> To calculate the global difference in 3′UTR isoform expression between each perturbation and the 97 samples containing non-targeting guide RNAs, the average difference in WUI (dWUI) was calculated. To do so, all multi-UTR genes with a mean expression of > 5 TPM and mean WUI-values between 0.1 and 0.9 in the samples containing the 97 non-targeting guide RNAs were analyzed and the difference in the mean WUI values between the perturbation and the control samples was calculated. Similarly, a difference in IPA (dIPA) values was calculated from IPA genes with a mean expression of > 5 TPM and mean IPA-values between 0.1 and 0.9 in cells receiving the non-targeting sgRNAs cells. For each perturbation, all average dWUI and dIPA values are reported in Table S6. These statistics average across all expressed multi-UTR or IPA genes for each perturbation. As a global average, it is most sensitive to unidirectional changes in WUI and IPA. Changes in opposite directions will cancel out.

<u>Clustering.</u> For each multi-UTR gene, we computed a baseline mean WUI value using the 97 non-targeting perturbations, weighted by the number of cells in each perturbation. We excluded genes that had more than 20% of cells non-detecting or a mean gene TPM lower than 20 in the non-targeting perturbations, leaving 1,775 genes. For all perturbations with at least 30 cells (*N* = 2,077), a dWUI was computed as a deviation from the baseline mean WUI. Missing WUI values in a target gene/perturbation pair were imputed as dWUI=0 (identical to baseline). A z-scaled dWUI (zdWUI) was computed by scaling the variance in dWUI across all targets to 1 without centering, since the center is already determined by the non-targeting perturbations. Three rounds of clustering were then performed using these zdWUI values.

In the first round of clustering, we identified and removed sets of perturbations that showed no pattern of regulation, which reduced the space from 2,077 genes to 30 principal components. Then, clustering was performed on the perturbations using walktrap community detection on a k=5 nearest neighbors graph. We examined the heatmaps of zdWUI of each perturbation cluster and observed that the two largest clusters had no visible patterns. Therefore, we removed all the 1,241 perturbations in these clusters from further analysis.

For the second round of clustering, we aimed to identify and remove sets of genes that showed no pattern of regulation. Dimensionality reduction and clustering was performed similar to before, but now on the perturbation space and with k=4. Three large clusters of genes showed no visual pattern of regulation, and their 892 genes were removed. The third round clustered the reduced set of 836 perturbations on the reduced set of 883 target genes. This final procedure used k=3 and identified 18 perturbation clusters and 17 target gene clusters.

<u>Identification of protein complexes and pathways within perturbation clusters.</u> To identify protein complexes or pathways within each of the 18 perturbation clusters, we generated a protein-protein interaction network based on the genes assigned to each perturbation cluster. Data from the String database v.11.5 were accessed through the StringApp and only high-confidence physical interactions (confidence score > 0.8) were included for the network creation. Figure 4b was generated using Cytoscape v.3.9.1, where average dWUI values of each perturbation were mapped to node colors and nodes were grouped by perturbation cluster. The dWUI values for each gene in each perturbation cluster are reported in Table S7.

<u>Differential gene expression and differential 3′UTR length within each perturbation cluster.</u> For dWUI testing, we required either the set of non-targeting perturbations or the perturbations in a given cluster to have a mean TPM > 5. We then performed a two-sided Mann-Whitney test between the WUI values for each gene comparing non-targeting and targeting perturbations. P-values were corrected for multiple testing using Benjamini-Hochberg procedure. Genes were considered significant if q-value < 0.05. Differential gene expression testing was similarly performed.

<u>Overlap of genes between perturbation clusters.</u> To find APA regulators that target similar genes, we determined the extent of overlap within their target genes. Gene overlaps between perturbation clusters were analyzed with the GeneOverlap R package v1.32.0^74^ and -log10-transformed p-values were reported (Fisher’s exact test).

<u>Classification of shortening or lengthening by isoform-specific regulation in each perturbation</u> <u>cluster.</u> For the genes with significant differential 3′UTR expression in each cluster (Fig. 5a), we analyzed isoform-specific regulation patterns. The analysis was limited to genes that had two 3′UTR isoforms.

First, we estimated isoform expression levels within a perturbation cluster from gene expression and WUI values with:

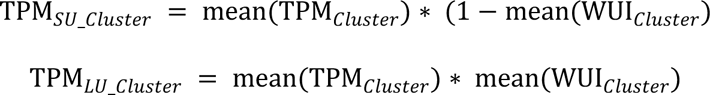

Next, cluster-specific absolute TPM changes (dTPM) relative to the control conditions were calculated for each isoform:

e.g.

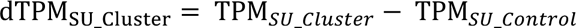

For compensatory (or coordinated) 3′UTR isoform regulation, we require that at least half of the expression gained by one isoform is lost by the other 3′UTR isoform in the gene, or vice versa:

Compensatory regulation, if:

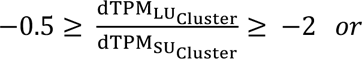

Conversely, genes for which this criterium was not met were categorized as having predominantly isoform-specific regulation, which was assigned to the isoform with the higher TPM expression difference (in absolute TPM values).

<u>Identification of gene features that correlate with dWUI in each perturbation cluster.</u> To investigate the relationship of particular gene features with 3′UTR isoform changes in each perturbation cluster we performed Pearson correlation. For all multi-UTR genes with a TPM > 5 in the non-targeting control condition, the mean dWUI value of each gene was correlated with their value for the respective gene feature. This analysis was performed for each perturbation cluster. For each feature, multiple testing correction was performed using Benjamini-Hochberg procedure and correlations with a q-value < 0.05 were considered significant.

Table S8 lists all correlation results, along with a comprehensive list of external data sources that were used for the analysis. When inferring 3′UTR lengths, we used the coordinate of the stop codon belonging to the longest annotated coding region of the gene. When inferring 3′UTR lengths and sequences, we used the coordinate of the stop codon belonging to the longest annotated coding sequence (CDS) of the gene (GENCODE v39). The same stop codon was assigned to all last exon 3′UTR isoforms of the gene and was also used for prediction of stop site readthrough (Table S1)^75^. Similarly, CDS and 5′UTR length, as well as other features related to their sequence, were determined based on mRNA annotations from the transcripts with the longest CDS. Features characterizing 3′UTR sequences, such as AU-rich elements, GC content and m6A modifications were assigned to both short and long 3′UTRs when occurring in the common region. For correlations related to splicing activity, a previously published dataset (Table S5)^76^ was used to identify exon junctions with the lowest predicted splice site score for the 5′ and 3′ splice site for each gene. For analyzing correlations regarding m6A methylation marks, previously identified m6A sites from a HeLa dataset^77^ were mapped to coding regions and 3′UTRs of all expressed multi-UTR genes expressed in K562 cells. Then, m6A scores were calculated as the sum of methylation levels at all sites within a region.

<u>Isoform-specific mRNA half-life analysis.</u> A K562 SLAM-seq experiment was used for mRNA stability analysis^78^. FASTQ files were processed using the SLAM-DUNK pipeline^79^ with our MWS UTRome as reference annotation. For each isoform, mean conversion rate per timepoint (0, 120, 240, 360 minutes) was computed as the weighted mean of conversions rates weighted by read counts (‘ReadsCPM’) across the three replicates. Mean conversion rates per isoform were modeled with first-order kinetics with the R ‘glm’ function, and mRNA half-life computed from the coefficients. The processing pipeline is available at https://github.com/Mayrlab/slam-k562-utrome.

## Data Availability

The following datasets were used:

Bulk 3′-seq datasets: HSC (ArrayExpress, E-MTAB-7391)^46^; ESC^49,50^ were obtained from the PolyASite v2.0 database^36^.

Mouse scRNA-seq datasets: Tabula muris (GEO:GSM3040890-917)^56^, bone marrow (GEO:GSM2877127-32)^47,48^, brain (GEO:GSM3722100-115)^57^, and ESC (GEO:GSM3629847-8)^51^, GSM4694997^52^.

Human scRNA-seq datasets: Tabula sapiens (https://tabula-sapiens-portal.ds.czbiohub.org)^55^, K562 Perturb-seq experiment (SRA:SRR19653800-847)^59^.

Perturb-seq feature comparison. Codon analysis and mRNA half-life measurements (GEO:GSE126522)^78^, m6A methylation (GEO:GSE211303)^77^.

## Code Availability

All code to generate figures is provided in the GitHub repository https://github.com/mayrlab/scUTRquant-figures. Code for data processing pipelines is available in the GitHub repositories listed with their respective methods. All code for scUTRquant, scUTRboot, and txcutr is open-source.

## Supplementary Figures

**Figure S1.**
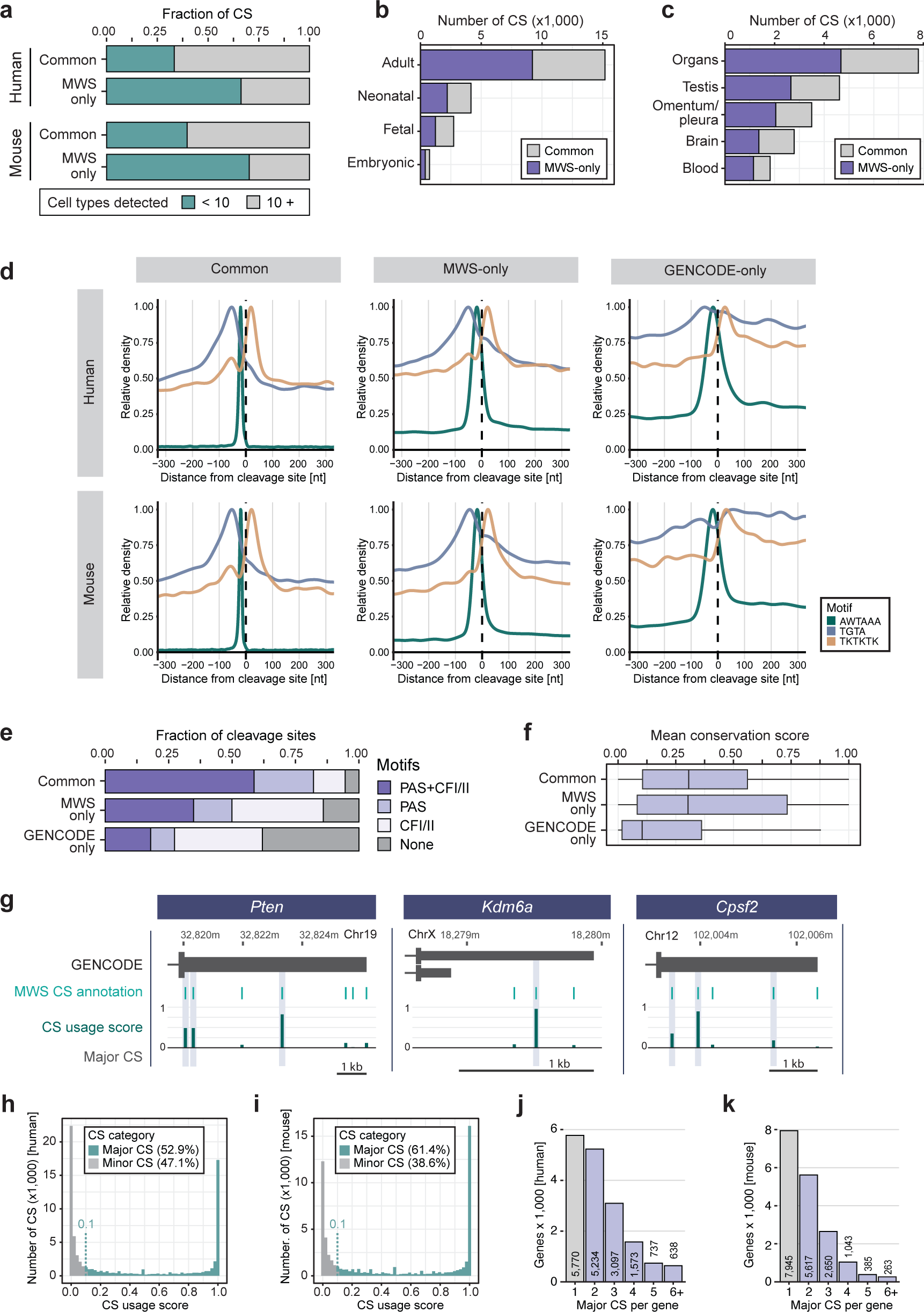
Characterization of mRNA 3′ end CS in 104 primary mouse cell types. **a,** Fraction of CS that were detected in fewer that ten cell types shown for common and novel CS. Shown are CS from human and mouse MWS annotations. **b,** For cell type-restricted mouse CS (detected in < 10 cell types), the developmental stage of the cell types is shown. **c,** As in **b**, but the tissue origin is shown. **d,** Metaplots for binding sites enrichment of CPA factors surrounding CS, including canonical PAS (AWTAAA, dark green), CFI (TGTA, blue) and CFII (TKTKTK, orange). **e,** As in **1c**, but binding site summary statistic is shown for mouse MWS data. **f,** As in **1e**, but PhastCons score was obtained from 60 species. **g,** As in **1f**, but example genes from the mouse transcriptome (mm10) are shown. **h,** CS usage score distribution for huma MWS data. **i,** As in **h**, but usage score distribution for mouse MWS data is shown. **j,** The number of major CS (obtained from Table S1) is shown for human genes that were analyzed by MWS. **k,** As in **j**, but for mouse genes (obtained from Table S2).

**Figure S2.**
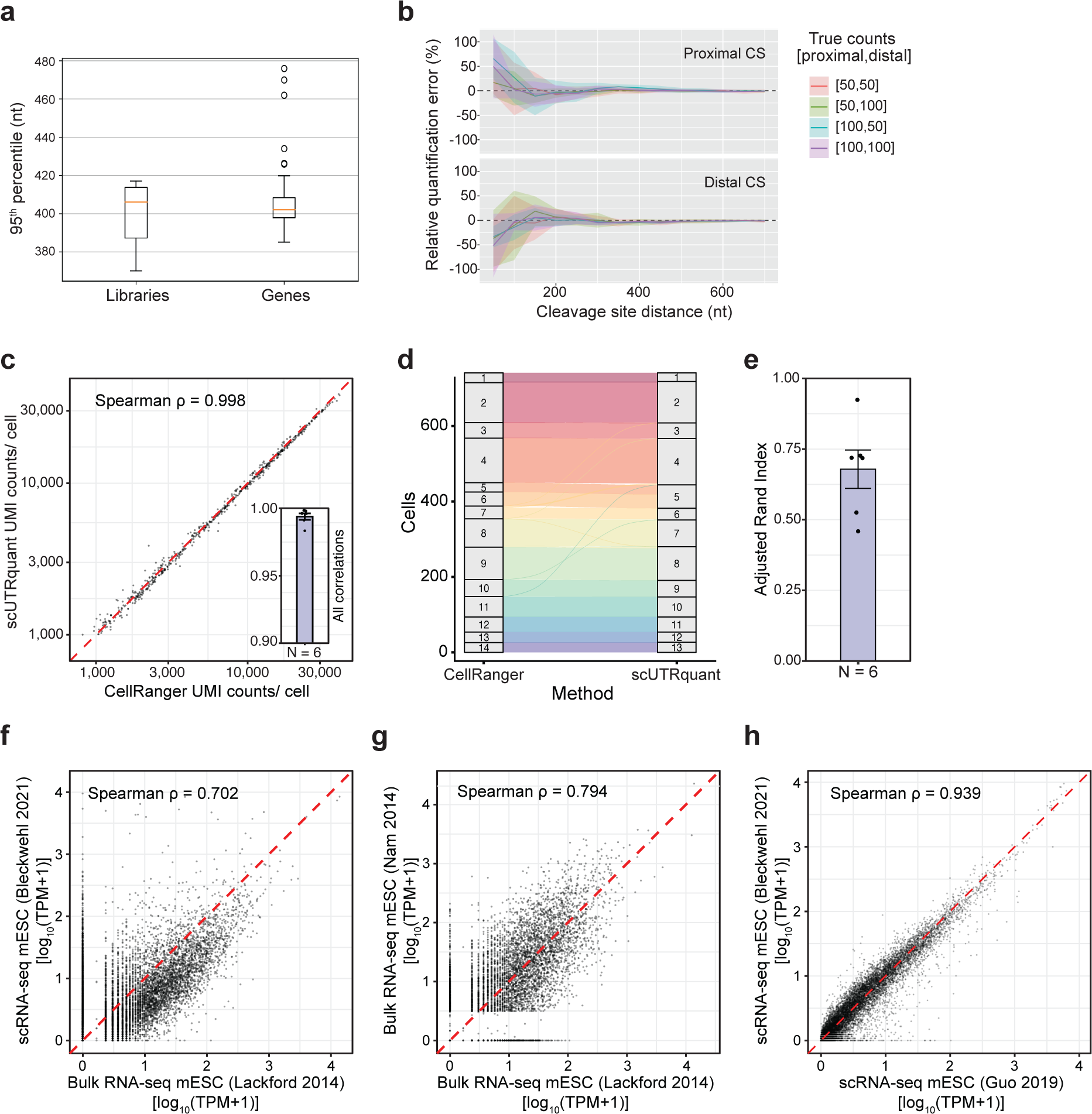
scUTRquant-derived gene and 3′UTR expression is precise and accurate. **a,** The cut-off for the UTRome truncation was empirically determined on mRNA 3′ ends for 56 curated genes (right) and 28 libraries from Tabula muris. The distance from the CS where the 95^th^ percentile read is located is shown as box plot. **b,** Error rates for transcript quantification with kallisto surveyed over a range of distances between CS. Lines indicate the mean of the relative errors from 10 simulations for each of four simulated input read configurations. Ribbons indicate two standard deviations from the mean. Mean relative errors stabilize with a minimum CS distance of 200 nt. **c,** Correlation of scUTRquant UMIs per cell obtained by scUTRquant and CellRanger for mouse heart 10x Genomics demonstration dataset. Inset: Additional Spearman correlations for six mouse 10x Genomics demonstration datasets, with mean and SE. **d,** Correspondence between clusters called from scUTRquant and CellRanger gene counts for the dataset in **c.** **e**, Adjusted Rand Indexes for six mouse 10x Genomics demonstration sets comparing clusters called from UMI counts from CellRanger versus scUTRquant. Mean and SE are shown. **f,** Correlation of 3′UTR isoform counts obtained by bulk 3′ end sequencing and scUTRquant isoform counts of scRNA-seq of mouse ESCs. **g,** Correlation of 3′UTR isoform counts between two experiments of bulk 3′ end sequencing for mouse ESCs. **h,** Correlation of scUTRquant 3’UTR isoform counts between two experiments of scRNA-seq of mouse ESCs.

**Figure S3.**
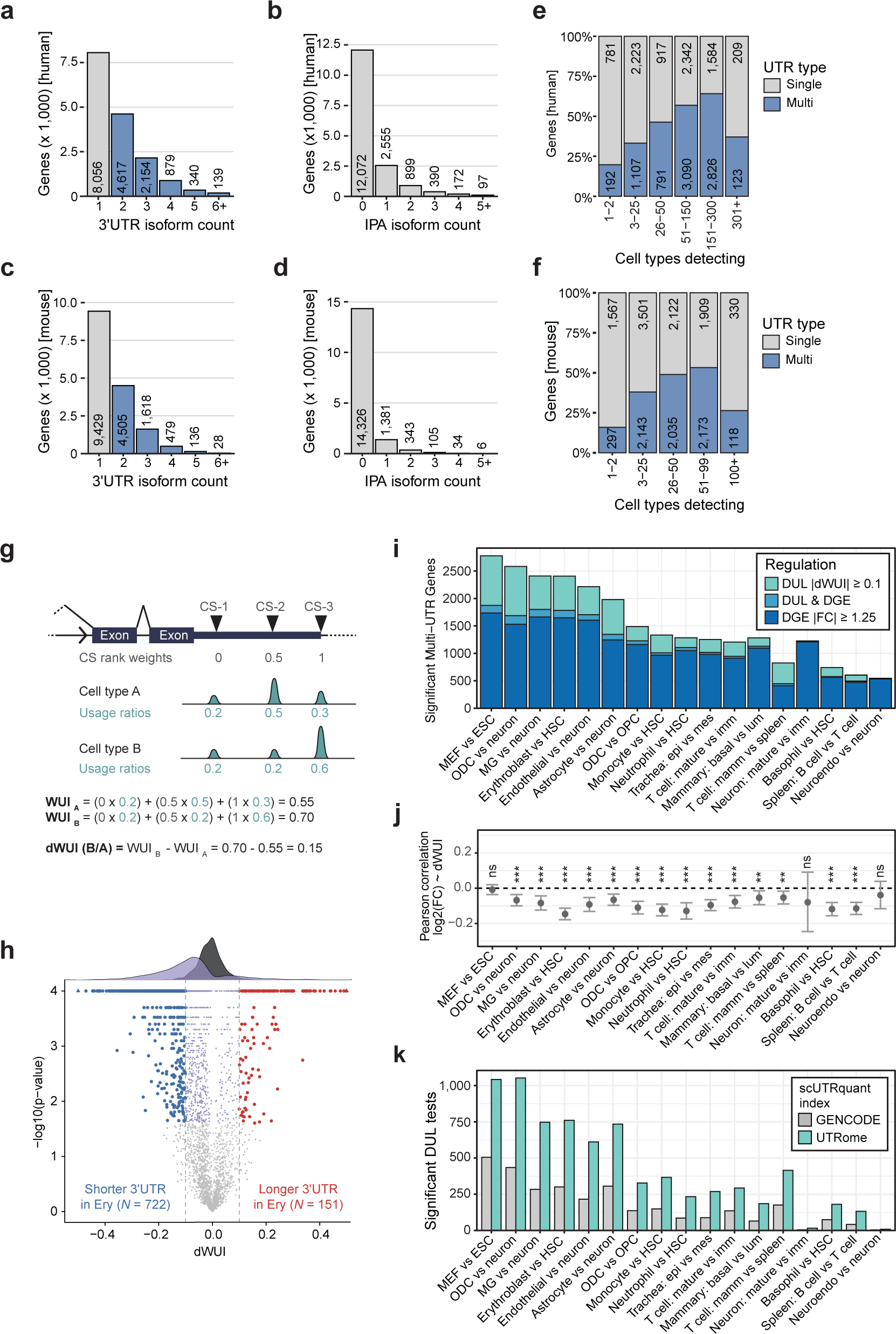
Application of scUTRquant to 355 human and 119 mouse cell types for comprehensive identification of single- and multi-UTR genes and genes with IPA isoforms. **a,** Distribution of single- and multi-UTR genes when using a minimum usage of 10% for each alternative 3’UTR isoform located in the terminal exon from 355 cell types obtained from the Tabula Sapiens data set. **b,** Same as **a**, but for IPA events. A minimum usage of 10% for each IPA isoform in any cell type. **c,** Same as **a**, but classified from a survey of 119 mouse cell types. **d,** Same as **b,** but classified from a survey of 119 mouse cell types. **e,** The fraction of single- and multi-UTR genes was plotted for human genes expressed broadly or in a cell type-restricted manner. Only genes detected in at least 50 cells were included. **f,** Same as **e**, but for shown are mouse genes. **g,** Example for calculation of Weighted UTR Index (WUI) for two hypothetical cell types and 3′UTR usage conditions. The change in WUI (dWUI) across conditions of 0.15 indicates a shift to expression of longer isoforms. **h,** Volcano plot showing p-values and the dWUI resulting from scUTRboot’s WUI bootstrap test between HSCs and Ery. Red, blue, significant dWUI (q-value < 0.05) and |dWUI| > 0.10. Purple, significant dWUI (q-value < 0.05), but with |dWUI| < 0.10 (*N* = 834); light grey, genes with non-significant dWUI (*N* = 1,693). **i**, As in **3f**, but with a 1.25 FC cutoff for gene expression change was applied. **j**, Pearson’s correlation coefficients for log2(FC) and WUI differences across 17 cell type comparisons shown in **i**. Error bars indicate 95% confidence intervals and significance of correlations are indicated with ns (not significant) if p-value > 0.05, ** p-value < 0.01, *** p-value < 0.001. **k,** As in **3g,** but shown is the number of genes with significant DUL when two different CS annotations were used. The number of significant DUL tests is shown across 17 comparisons of mouse cell types using our UTRome compared with CS annotations from GENCODE.

**Figure S4.**
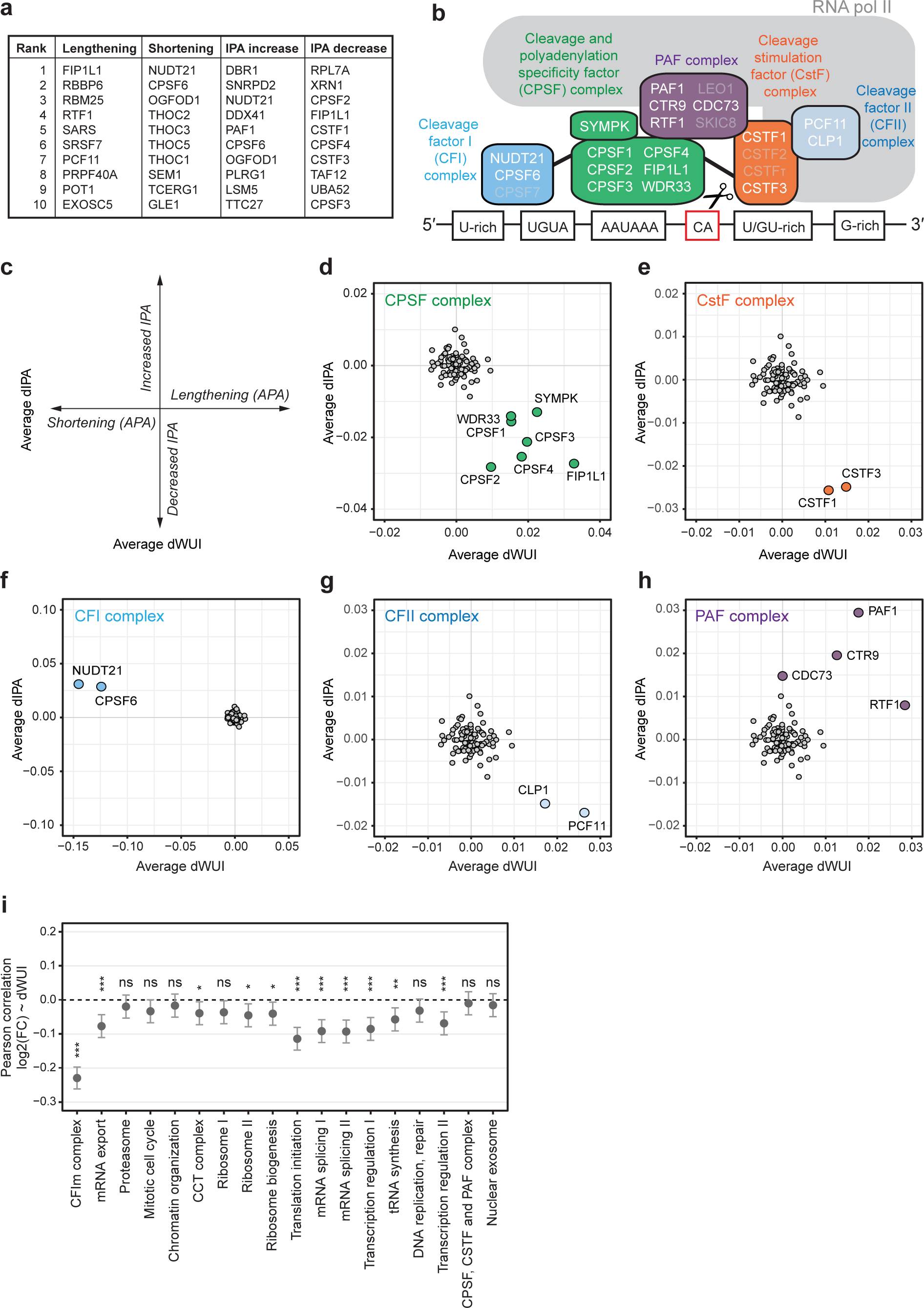
scUTRquant identifies known APA and IPA regulators in a Perturb-Seq data set. **a,** Ranked list of top ten regulators of APA and IPA among essential perturbations in K562 cells. A full list of all summarized perturbation effects is provided in Table S6. **b,** Schematic representation of the CPA machinery. Genes listed in light grey are not present in the data set. **c,** Plotting of global shifts in APA and IPA isoform expression patterns. **d,** Knockdown effects of genes constituting the CPSF complex (green) as shown in **b**, and non-targeting control guide RNAs (grey, *N* = 97). **e,** As in **d**, but with components of the CSTF complex (orange). **f,** As in **d**, but with components of the CFI complex (blue). **g,** As in **d**, but with components of the CFII complex (light blue). **h,** As in **d**, but with components of the PAF complex (purple). **i,** Pearson’s correlation coefficients for log2(FC) and WUI differences across the 18 perturbation clusters from **4c**. Error bars indicate 95% confidence intervals and significance of correlations are indicated with ns (not significant) if q-value > 0.05; ** q-value < 0.01, *** q-value < 0.001.

**Figure S5.**
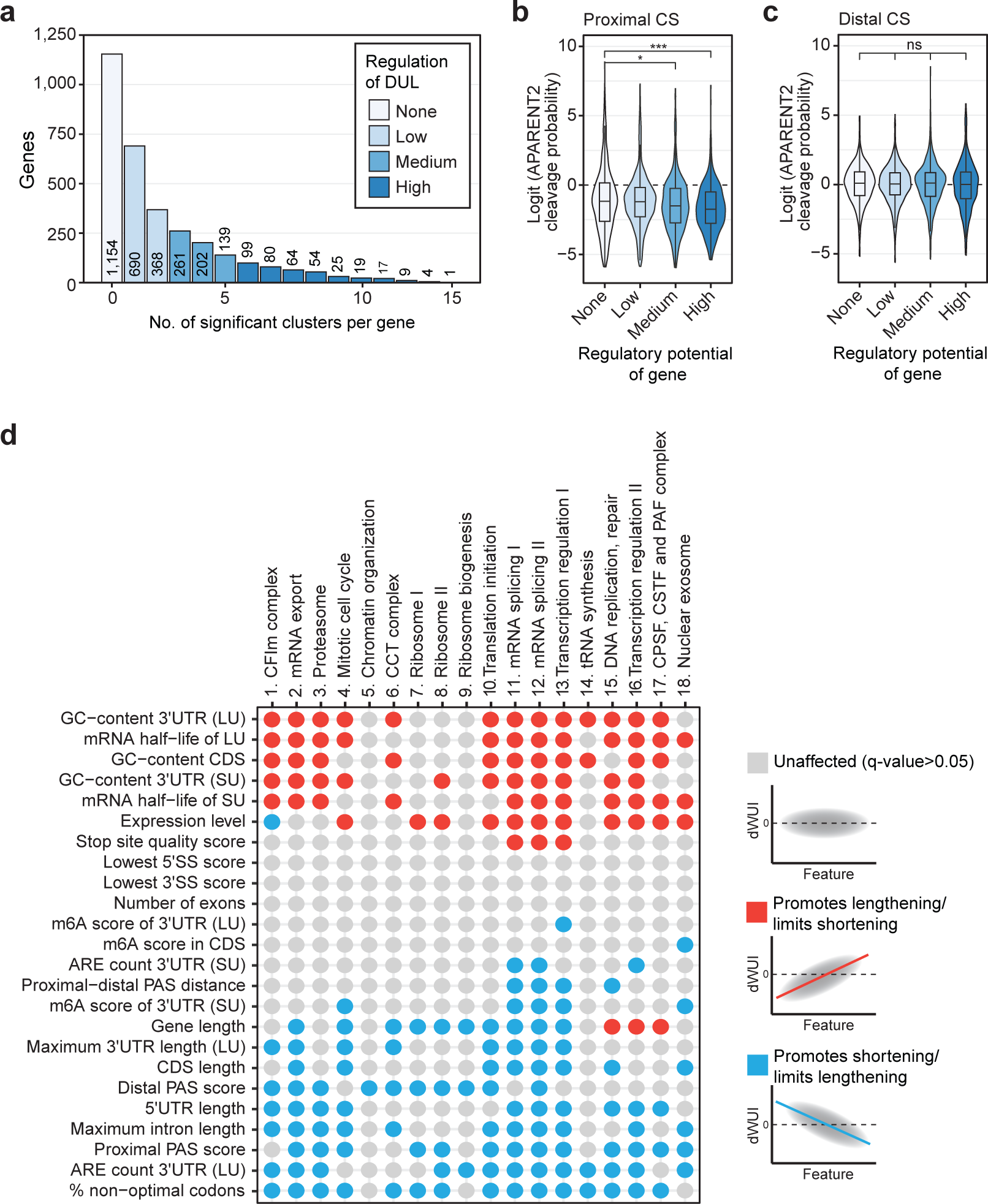
The regulatory potential with respect to APA is mostly determined by the proximal CS. **a,** Distribution plot of multi-UTR genes showing the number of perturbation clusters that cause significant 3′UTR isoform expression changes. Genes were categorized into one of four groups based on their potential for dynamic regulation. **b,** Analysis of predicted CS efficiencies across gene groups exhibiting different regulatory potential for proximal CS. APARENT2 cleavage probability scores are used as measure for CS strength. Box shows IQR with median and whiskers 1.5*IQR. Statistical significance was calculated using a two-sided Mann-Whitney test (n.s. not significant, * *P* < 0.05, *** *P* < 0.001). **c,** As in b, but shown is APARENT2 cleavage probability scores for distal CS. **d,** Gene and mRNA features that correlate with changes in 3′UTR isoform expression. Summarized are the results of correlations between each feature and each perturbation cluster. Grey dots represent correlations with FDR-corrected p-values > 0.05, while red and blue dots mark significant positive or negative correlations between features and average dWUI values in each perturbation cluster.

## Notes

https://github.com/Mayrlab/scUTRquant

https://github.com/Mayrlab/scUTRquant-figures

### Competing Interest Statement

The authors have declared no competing interest.

### Summary of Updates

The annotation pipeline was reworked to identify cleavage sites at the celltype level, rather than in aggregate. Annotation is also now added for human. Several quality checks are added for the novel cleavage sites. All previous analyses are now run with the updated annotations. New analyses of Tabula Sapiens and a Perturb-seq dataset are added, including extensive characterization of perturbations that impact UTR isoforms. LUI statisitics are replaced with WUI statistics. Sibylle Mitschka is added as author who worked on Perturb-seq analysis; Gang Zhen is removed as his contribution from the previous work is no longer included.

